# Cyclin B3-CDK1 couples translational de-repression to embryonic mitoses via OMA protein degradation in *C. elegans*

**DOI:** 10.64898/2026.07.22.740101

**Authors:** David Bojorquez, Shabnam Moghareh, Maab Elsayed, Amrutha Kizhedathu, Gio Jison, Lee Bardwell, Pablo Lara-Gonzalez

**Author notes:** Corresponding author: Email for manuscript correspondence, Address: University of California Irvine, McGaugh Hall 4244, Irvine CA 92697.

## Abstract

During early embryogenesis, gene expression relies on maternally loaded mRNAs whose translation is controlled by RNA binding proteins. Here, we identify a critical role for the cyclin B3-CDK1 complex, known for its function in mitosis, in driving early embryonic gene expression in *C. elegans*. The cyclin B3-CDK1 complex works by marking the RNA binding OMA proteins (OMA-1 and OMA-2) for degradation, which ensures the de-repression and translation of their target mRNAs. OMA protein degradation relies on cyclin B3’s conserved phosphate-binding pocket, which promotes multi-site OMA phosphorylation and the generation of phospho-degrons. Notably, the phosphate-binding pocket of cyclin B3 does not substantially contribute to its mitotic roles, indicating that the mitotic and translational functions of the cyclin B3-CDK1 complex are separable. These findings establish that embryonic activation of the cyclin B3-CDK1 complex drives both mitotic divisions and mRNA de-repression, which ensures that cell division is coupled to the early gene expression program in development.

## INTRODUCTION

The success of embryonic development depends on the coordination of cell division with cell fate specification and differentiation. Following fertilization, embryos undergo a series of rapid divisions characterized by the absence of gap phases and consisting primarily of rounds DNA replication followed by mitosis. Concurrent with this process, the expression of maternally loaded mRNAs and subsequent activation of the zygotic genome drives cells to acquire distinct fates to ultimately give rise to the specialized tissues of the developing embryo (Shahbazi, 2020; Verlhac et al., 2010). Defining the molecular mechanisms that link mitotic divisions to cell fate specification is essential to understanding embryonic development.

The *Caenorhabditis elegans* embryo is a powerful model system for studying early developmental events, due to its rapid and invariant embryonic development, combined with a fully mapped cell lineage, optical transparency, and a robust genetic toolkit (Robertson and Lin, 2015). After fertilization, *C. elegans* embryos undergo several rounds of asymmetric cell division to give rise to blastomeres with defined fates (Rose and Gonczy, 2014). Central to this process is the regulation of maternally deposited transcripts, which are initially silenced and later segregated and expressed to drive early cell fate specification. The <u>O</u>ocyte <u>MA</u>turation proteins OMA-1 and OMA-2 (Detwiler et al., 2001), two RNA binding proteins that share homology with the mammalian Zinc Finger Protein 36 (ZFP36) family of proteins, are essential regulators of early mRNA expression. As they accumulate in maturing oocytes, OMA proteins bind to the 3’UTR of their target mRNAs via a triple arginine motif and a CCCH-type tandem zinc finger domain that recognizes repetitive UA(AU) motifs (Ertekin et al., 2025; Kaymak and Ryder, 2013). Immunopurification and ribosome profiling experiments have identified thousands of early expressed maternal transcripts as targets of OMA, including *zif-1, mom-2, nos-2 and cdc-25.3* (Oldenbroek et al., 2013; Shukla et al., 2025; Spike et al., 2014; Tsukamoto et al., 2017). Binding of OMA proteins to their target mRNAs ensures their translational repression, although OMAs can also promote translation for specific transcripts such as *spn-4* and *meg-1* (Tsukamoto et al., 2017). In addition to mRNA repression, OMA proteins have also been proposed to suppress RNA polymerase II – dependent transcription in zygotes by binding to the transcription initiation factor TAF-4 and sequestering it in the cytoplasm (Guven-Ozkan et al., 2008). Importantly, upon fertilization, OMA-1 and OMA-2 rapidly degrade through the action of several E3 ubiquitin ligases of the Cullin-RING family (Du et al., 2015). Timely degradation of OMA proteins is critical for the de-repression of maternally loaded transcripts, placing it as an essential event in the initiation of transcript expression in the early embryo. Consistent with this, mutations in OMA that impair its degradation cause severe disruptions to early cell fate specification, resulting in embryonic lethality (Lin, 2003). However, despite the critical role of OMA degradation in embryonic development, the precise mechanisms by which OMA is degraded upon fertilization remains poorly understood.

Genetic screens have identified several protein kinases that participate in OMA protein degradation (Nishi and Lin, 2005; Shirayama et al., 2006). These include the Dual Specificity Tyrosine-Phosphorylation-Regulated Kinase 2 (DYRK2) orthologue MBK-2, Glycogen Synthase Kinase 3 (GSK-3) and the Casein Kinase 1 orthologue KIN-19 (Nishi and Lin, 2005; Shirayama et al., 2006). Of these MBK-2, which activates shortly after embryo fertilization, is essential for OMA degradation by phosphorylating a key residue in the unstructured C-terminus of OMAs (Nishi and Lin, 2005). Preventing phosphorylation at OMA-1’s MBK-2 site (Thr239) results in its increased stability and prolongs the repression of OMA mRNA targets, resulting in penetrant embryonic lethality (Lin, 2003). Interestingly, the mitotic kinase CDK-1 was found to be important for OMA degradation. During mitosis, CDK-1 is activated by cyclin B to orchestrate several mitotic events, such as nuclear envelope breakdown, chromosome alignment, spindle assembly and chromosome segregation, among others (Crncec and Hochegger, 2019; Lindqvist et al., 2009). Cyclin B–CDK-1 complexes phosphorylate target substrates on consensus [S/T]-P or [S/T]-P-x-[K/R] sequences to promote timely mitotic progression (Crncec and Hochegger, 2019; Lindqvist et al., 2009). Most metazoans encode for three isoforms of cyclin B that activate Cdk1: cyclin B1 (CYB-1), cyclin B2 (CYB-2) and cyclin B3 (CYB-3). While CYB-1 and CYB-2 are closely related in sequence and function, CYB-3 diverges and shares similarity to A-type cyclins (Gallant and Nigg, 1994). Notably, CYB-3 is a dominant activator of CDK-1 in early embryonic mitoses in *C. elegans* (Deyter et al., 2010; Lara-Gonzalez et al., 2024; van der Voet et al., 2009) and has also been implicated in meiotic progression in vertebrate oocytes, including mammals (Fatemi et al., 2021; Karasu et al., 2019; Li et al., 2019; Zhu et al., 2026). In addition to activating CDK-1, cyclin B proteins provide substrate specificity by directly recognizing Short Linear Motifs (SLiMs) in CDK-1 substrates. This recognition is mediated through two substrate binding surfaces: the hydrophobic patch (HP) and the phosphate binding pocket (PBP) (Loog and Morgan, 2005; Yu et al., 2021). While the HP is defined by a conserved MRAIL ([M]-[R]-x-[I]-[L/V]-x-x-[W]) sequence that recognizes the “RxL” SLiM in target substrates (Loog and Morgan, 2005), the PBP harbors a basic patch that directly interacts with pre-phosphorylated substrates (Yu et al., 2021). Together, these substrate binding surfaces tether the cyclin B–CDK-1 complex to its substrates, facilitating their phosphorylation (Asfaha et al., 2022; Heinzle et al., 2025; Loog and Morgan, 2005; Ng et al., 2025; Schunk et al., 2025). Although CDK-1 inhibition delays OMA degradation (Shirayama et al., 2006), whether this was an indirect consequence of impaired mitotic progression was unclear.

In this study we demonstrate that the CYB-3–CDK-1 complex, acting through CYB-3’s conserved phosphate binding pocket (PBP), is a key mediator of OMA degradation *in vivo*. Mutating CYB-3’s PBP resulted in severe OMA stabilization and embryonic lethality, but it did not substantially affect CYB-3–CDK-1’s ability to promote mitotic progression, thus decoupling CDK-1’s roles in development from its roles in mitosis. Using a combination of OMA degradation reporters and *in vitro* biochemical reconstitution experiments we identified a novel site on OMA that is crucial for CYB-3–CDK-1 – dependent OMA hyper-phosphorylation *in vitro* and OMA degradation *in vivo*. Together, our findings support a model in which the CYB-3–CDK-1 promotes the degradation of OMA proteins during early embryogenesis via CYB-3’s PBP. We propose a ‘saturation model’ of phosphorylation, in which multiple kinases collectively drive timely OMA degradation by phosphorylating a combination of critical and redundant sites, which in turns ensures normal embryonic patterning.

## RESULTS

### OMA protein degradation begins during pronuclear meeting

In *C. elegans*, the degradation of OMA proteins is an essential event that promotes the de-repression of maternal mRNAs to drive early embryo mRNA translation (**Fig. 1A**). Although OMA proteins have been shown to degrade sometime between the one-and two-cell stage of embryonic development, the precise timing of this degradation has not been reported. Early *C. elegans* embryogenesis is defined by a series of coordinated events, which include the establishment of anterior-posterior polarity, the meeting of the maternal and paternal pronuclei, entry into the first mitotic division, and the subsequent asymmetric cleavage that gives rise to daughter cells of distinct developmental fates (Oegema and Hyman, 2006).

**Figure 1.**
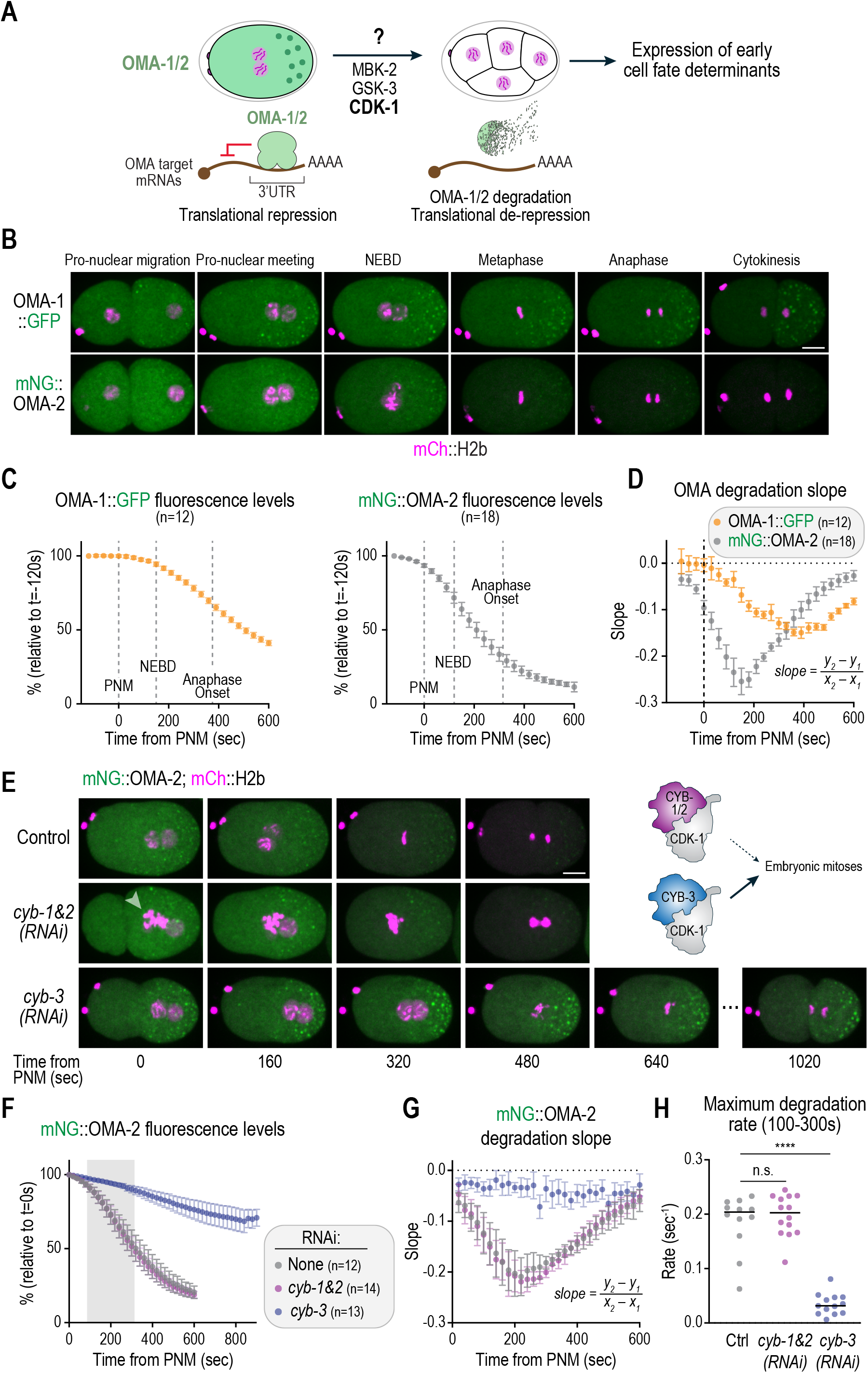
OMA proteins degrade at pro-nuclear meeting in a CYB-3–CDK-1 – dependent manner. **(A)** Schematic illustrating how OMA-1/2 degradation promotes maternal mRNA translation and early cell fate specification in *C. elegans* embryos. Multiple kinases, including MBK-2, GSK-3 and CDK-1, participate in ensuring OMA protein degradation in early embryogenesis. **(B)** Stills from time-lapse sequences of embryos expressing mCherry-tagged Histone H2b, and either OMA-1::GFP or mNeonGreen::OMA-2, undergoing pronuclear migration and their first embryonic division. **(C)** Quantification of OMA-1::GFP and mNG::OMA-2 fluorescence intensity signals over time for the conditions in *(B)*. The average timings for nuclear envelope breakdown (NEBD) and anaphase onset, relative to pronuclear meeting, are shown. **(D)** Second derivative analysis of the data in *(B)*. **(E)** *(left)* Stills from time-lapse sequences of embryos expressing mCherry-tagged Histone H2b and mNeonGreen-tagged OMA-2, imaged under the specified conditions. Arrowhead indicates a maternal pronucleus with extra chromosomes, as it is characteristic for *cyb-1&2(RNAi)* embryos (Lara-Gonzalez et al., 2024). *(right)* Schematics illustrating the role of cyclin B-CDK-1 complexes in promoting embryonic mitoses in *C. elegans.* **(F)** Quantification of mNG::OMA-2 fluorescence intensity levels for the conditions in *(E)*. Gray zone represents time of maximum OMA degradation, used for calculating degradation rates in *(H)*. **(G)** Second derivative analyses for the data in *(F)*. **(H)** Quantification of maximum degradation rate from slopes between 100-300 sec post-PNM from *(F) (See also Fig. S1F)*. *n* represents the number of embryos analyzed. Scale bars, 10 µm. Error bars are 95% intervals. **** represents *P* < 0.0001 from Mann-Whitney tests; non-significant (n.s.) is *P* > 0.05.

To define OMA degradation kinetics in early embryogenesis, we developed a time-lapse fluorescence microscopy assay. For this, we took advantage of two previously published *C. elegans* lines in which OMA-1 and OMA-2 are endogenously tagged with the fluorescent proteins GFP and mNeonGreen, respectively. Both OMA fusions are functional, as each is individually capable of supporting oocyte development in the absence of the other (Ertekin et al., 2025; Hu et al., 2021) (**Fig. S1A&B**). Using these fluorescently tagged OMA lines, we found that degradation of both OMA-1 and OMA-2 initiates at around the time of the meeting of the maternal and paternal pronuclei, referred to as pronuclear meeting (PNM) (**Fig. 1B& C; Movie S1**). Second derivative analyses revealed that, while OMA-2 degradation begins shortly before PNM, OMA-1 degradation appears to start 60-90 sec post-PNM and proceeds through early embryogenesis with slower kinetics than OMA-2 (**Fig. 1D**). Taken together, these data indicate that OMA degradation is initiated at the time of pronuclear meeting, prior to the embryo’s commitment to the first mitotic division, which is substantially earlier than previously appreciated. The faster degradation kinetics of OMA-2 make it more amenable to quantitative analysis; therefore, we focused on OMA-2 for all subsequent experiments.

### CYB-3–CDK-1 activity is essential for timely OMA degradation

As stated above, several protein kinases have been implicated in promoting OMA protein degradation, including MBK-2 (DYRK2 in vertebrates), GSK-3 and CDK-1 (Nishi and Lin, 2005; Shirayama et al., 2006) (**Fig. 1A**). While the contributions of MBK-2 and GSK-3 have been defined in earlier work (Nishi and Lin, 2005), how CDK-1 fits into this process remains an open question. To begin addressing this, we first re-evaluated the specific requirement for individual cyclin B isoforms within the cyclin B-CDK-1 complex (Shirayama et al., 2006). In *C. elegans* embryos, cyclin B1 (CYB-1), cyclin B2 (CYB-2), and cyclin B3 (CYB-3) are redundantly required for the initiation of mitosis, with cyclin B3 playing a critical role as a hyperactivator of CDK-1 that accelerates the pace of embryonic mitoses (Lara-Gonzalez et al., 2024; van der Voet et al., 2009). Using RNA interference (RNAi), we found that co-depletion of CYB-1 and CYB-2 (referred to herein as *cyb-1&2(RNAi)*) had no significant effect on OMA-2 degradation kinetics (**Fig. 1E-G**). However, depletion of CYB-3 severely delayed OMA-2 degradation (**Fig. 1E-G**), which is consistent with prior work evaluating OMA-1 degradation through end point analyses (Shirayama et al., 2006). We validated the efficiency of our RNAi depletions through the recapitulation of previously reported mitotic phenotypes associated with CYB-1&2 versus CYB-3 depletions(Lara-Gonzalez et al., 2024) (**Fig. S1C-E**). To quantify this delay, we calculated the maximum degradation rate by measuring the slope of the degradation curves between 100-300 sec post-PNM (**Fig. S1F**), which indicated that CYB-3 depletion reduced the OMA-2 degradation rate to ∼19% of controls (**Fig. 1H**). These findings establish that CYB-3–CDK-1 activity is specifically required for OMA degradation following pronuclear meeting.

### OMA degradation requires CYB-3’s phosphate binding pocket

Having confirmed that CYB-3–CDK-1 activity is required for OMA degradation, we next set out to explore the underlying mechanism. For this, we turned our attention to CYB-3’s substrate binding surfaces, namely the hydrophobic patch (HP) and the phosphate binding pocket (PBP) (Loog and Morgan, 2005; Yu et al., 2021) (**Fig. 2A**). To test if putative substrate binding motifs within CYB-3 contribute to OMA degradation, we leveraged our previously established CYB-3 replacement system (Lara-Gonzalez et al., 2024). We engineered constructs encoding for RNAi-resistant CYB-3 transgenes where we mutated key residues within the hydrophobic patch (T143A, L147A, W150A; HP^mut^) and the phosphate binding pocket (R250A, R267A; PBP^mut^) (**Fig. 2A**). Both CYB-3 HP^mut^ and PBP^mut^ mutants were expressed at levels comparable to wild-type CYB-3 (**Fig. 2B**). In addition, both CYB-3 HP^mut^ and PBP^mut^ retained their ability to co-immunoprecipitate with CDK-1 when expressed as mNeonGreen fusion proteins, whereas a CDK-1 binding mutant (Y112A, I117A, Y119A; CB^mut^) did not (**Fig. 2C**). Notably, mutating CYB-3’s PBP substantially delayed OMA-2 degradation, to a similar extent as CYB-3 depletion (**Fig. 2D-F; Movie S2**). On the other hand, mutating CYB-3’s HP resulted in a much less severe delay in OMA-2 degradation (**Fig. 2D-F; S1C-E; Movie S2**).

**Figure 2.**
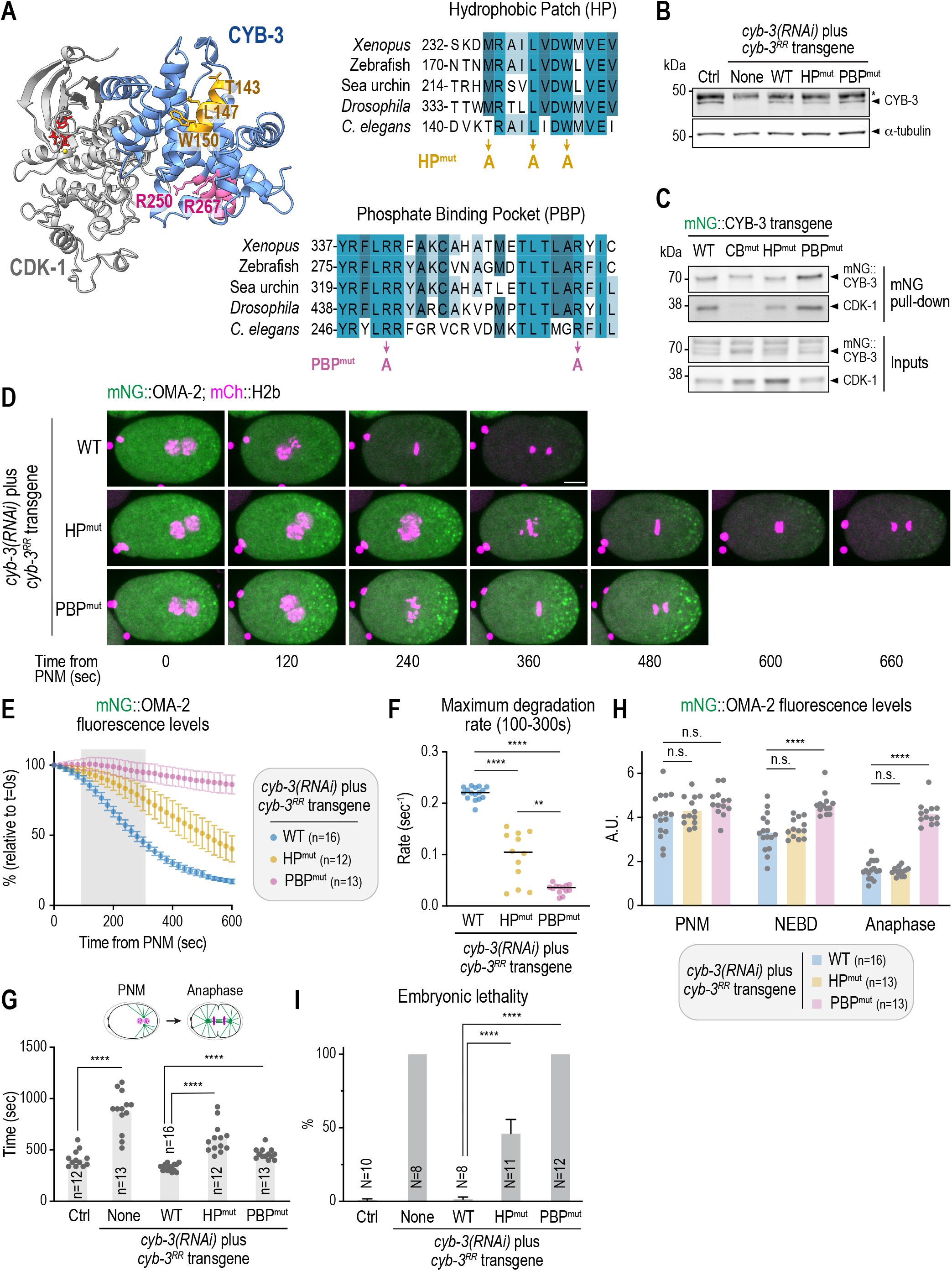
OMA-2 degradation is driven by CYB-3’s phosphate binding pocket (PBP). **(A)** *(left)* AlphaFold3 model of CDK-1 (grey) in complex with CYB-3 (blue). The hydrophobic patch (HP; yellow) and the phosphate binding pocket (PBP, magenta), as well as ATP (red) are shown. *(right)* Alignment of cyclin B3 sequences from different species illustrating conservation of substrate binding surfaces across evolution. The residues mutated to generate HP^mut^ and PBP^mut^ versions of CYB-3 are shown. **(B)** Immunoblots of *C. elegans* extracts showing expression of CYB-3 transgenes. α-Tubulin serves as a loading control. The asterisk denotes an unspecific band detected by the anti-CYB-3 antibody. **(C)** Immunoblot showing immunoprecipitation of mNG-tagged CYB-3 (WT, CB^mut^, HP^mut^ or PBP^mut^) from whole worm lysates. CB^mut^ corresponds to mutant deficient in CDK-1 binding (Lara-Gonzalez et al., 2024) and it is used as a negative control. **(D)** Stills from time-lapse sequences of embryos expressing mCherry-tagged Histone H2b, mNeonGreen-tagged OMA-2 and the indicated *cyb-3* transgenes, undergoing early embryogenesis. Scale bar, 10 µm. **(E)** Quantification of mNG::OMA-2 fluorescence intensity levels for the conditions illustrated in *(D)*. Gray zone represents time of maximum OMA degradation, used for calculating degradation rates in *(F)*. Error bars are 95% confidence intervals. **(F)** Quantification of maximum degradation rate from slopes between 100-300 sec post-PNM from *(E)*. **(G)** Quantification of the timing from PNM to Anaphase onset for the indicated conditions. These data are the same as for *Fig. S1E.* **(H)** Quantification of OMA-2 levels at PNM, NEBD and Anaphase for the specified conditions. **(I)** Quantification of embryonic lethality in animals depleted of CYB-3 through RNAi and expressing the indicated *cyb-3* transgenes. *n* represents the number of embryos analyzed; *N* represents the number of adults whose progeny was scored. **** represents *P* < 0.0001 from Mann-Whitney tests; * represents *P* < 0.05; non-significant (n.s.) is *P* > 0.05.

During our OMA degradation analyses we observed that, compared to wild-type CYB-3-expressing embryos, CYB-3 PBP^mut^ embryos displayed an ∼133 sec delay in the timing from PNM to anaphase (**Fig. 2G**). On the other hand, CYB-3 HP^mut^ embryos showed a longer delay of ∼280 sec in the PNM to Anaphase transition. Both CYB-3 mutant delays were significantly shorter than the ∼460 sec delay observed upon CYB-3 depletion (**Fig. 2G**). To address whether overall cell cycle delays account for the delays in OMA-2 degradation in CYB-3 HP and PBP mutants, we measured OMA-2 levels at PNM, NEBD and Anaphase onset. Strikingly, CYB-3 PBP^mut^ embryos showed impaired OMA-2 degradation even after correcting for cell cycle stage (**Fig. 2H**), indicating that the effect of CYB-3 PBP^mut^ in OMA degradation is independent of cell cycle progression. In contrast, CYB-3 HP^mut^ embryos had similarly reduced OMA-2 levels at PNM, NEBD and Anaphase, suggesting that the delay in OMA-2 degradation in CYB-3 HP^mut^ can be attributed to reduced CDK-1 activity and slower cell cycle progression. We conclude that the CYB-3–CDK-1 complex drives OMA degradation through substrate recognition via CYB-3’s phosphate binding pocket.

### Cyclin B3’s phosphate binding pocket is largely dispensible for mitotic progression

While CYB-3 depletion resulted in ∼3 fold mitotic delay and an increase in the rate of chromosome segregation errors, CYB-3 PBP^mut^ embryos showed no apparent mitotic defects and exhibited a milder (∼80 sec) mitotic delay (**Fig. S1D&G**). These data suggest that the CYB-3 PBP does not majorly contribute to mitotic events. Yet, despite its relatively modest mitotic phenotype, CYB-3 PBP^mut^ embryos exhibit 100% embryonic lethality (**Fig. 2I**). On the other hand, CYB-3 HP^mut^ embryos exhibited a longer mitotic delay than CYB-3 PBP^mut^ embryos (**Fig. S1D**) but had lower embryonic lethality (**Fig. 2I**), consistent with a partial loss-of-function phenotype. Taken together, these observations strongly suggest that the lethality associated with CYB-3 PBP^mut^ is driven primarily by developmental defects caused at least in part by impaired OMA degradation, rather than by mitotic errors.

### Cyclin B3’s phosphate binding pocket is critical for the translation of OMA mRNA targets

As described above, OMA proteins function in the early embryo by repressing the translation of maternally loaded mRNAs, and timely OMA degradation is required for translational de-repression (Oldenbroek et al., 2013; Shukla et al., 2025; Spike et al., 2014; Tsukamoto et al., 2017). To evaluate if the CYB-3 PBP is required for 3’UTR-driven mRNA translation in the early embryo, we generated translational reporter transgenes for established OMA target transcripts by fusing their 3’UTR to StayGold2::Histone H2b, under a germline promoter (*Pmex-5*) (Guven-Ozkan et al., 2010; Oldenbroek et al., 2013) (**Fig. 3A**). We selected two mRNAs, *zif-1* and *mom-2*, based on their previously reported high binding to OMA proteins and their characteristic de-repression at the 4 to 8 cell stage (Guven-Ozkan et al., 2010; Oldenbroek et al., 2013) (**Fig. 3A**). As previously reported, we found that in wild-type CYB-3 embryos, *zif-1* and *mom-2* are robustly translated in the AB and P4 lineages of embryonic development, respectively (**Fig. 3B-E; S2**). Importantly, mutating the PBP in CYB-3 severely diminished the expression of *zif-1* and *mom-2* translational reporters (**Fig. 3B-E**). These data are consistent with the CYB-3–CDK-1 complex being essential for the degradation of OMA proteins and the de-repression of OMA target mRNAs in early embryogenesis.

**Figure 3.**
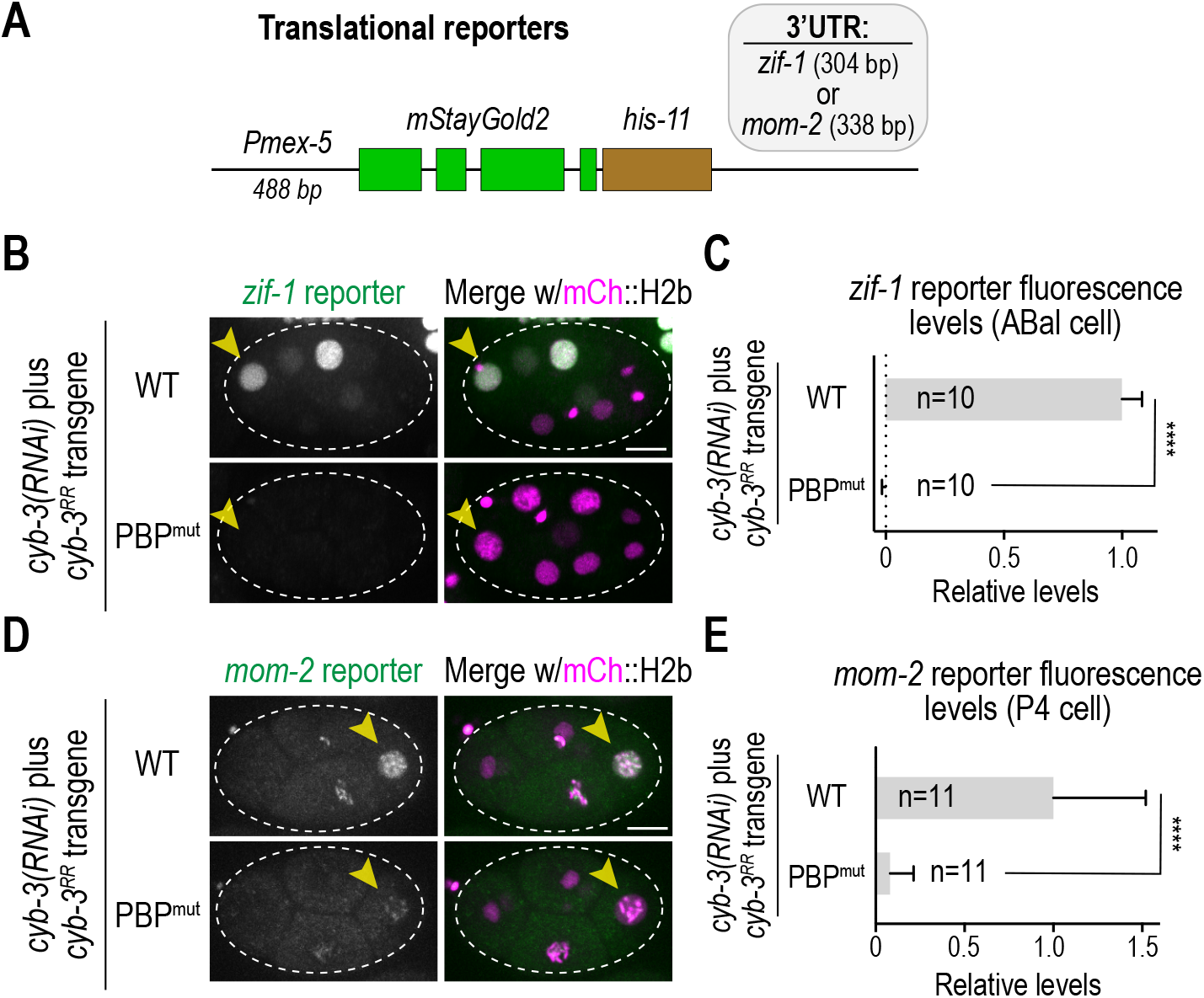
The CYB-3 PBP is crucial for the de-repression of *zif-1* and *mom-2* transcripts. **(A)** Schematic illustrating the translation reporters engineered to quantify *mom-2* and *zif-1* transcript de-repression. Note that *mom-2* and *zif-1* reporters only differ in their 3’UTR sequence but are otherwise identical. **(B)&(D)** Stills from time-lapse sequences of embryos expressing mCherry-tagged Histone H2B, and either *zif-1 (B)* or *mom-2 (D)* translational reporters, in addition to the corresponding *cyb-3* transgenes. Arrowheads indicate the ABal *(B)* or P2 *(D)* precursor. **(C)&(E)** Quantification of *zif-1 (C)* and mom-2 *(E)* translational reporter fluorescence intensity levels in the ABal *(C)* and P2 *(E)* precursor, under the specified conditions. *n* represents the number of embryos analyzed. Scale bars, 10 µm. Error bars are 95% intervals. **** represents *P* < 0.0001 from Mann-Whitney tests; non-significant (n.s.) is *P* > 0.05. *See also Figure S2*.

### The cyclin B3 phosphate binding pocket promotes OMA hyperphosphorylation

So far, our findings revealed that the phosphate binding pocket of CYB-3 is essential for timely OMA degradation and the de-repression of OMA mRNA targets in the early *C. elegans* embryo. We next sought to understand the molecular mechanism by which the CYB-3 PBP promotes OMA degradation. For this, we first aimed to define the domain in OMA that is sufficient for its degradation. OMA proteins possess a structured N-terminal half consisting of two predicted alpha helix domains as well as a Zn^2+^ finger domain (ZFD) that directly interacts with mRNAs’ 3’UTR (Detwiler et al., 2001). In addition, OMA proteins have a predicted intrinsically disordered C-terminal domain that shares ∼72% similarity between OMA-1 and OMA-2 (**Fig. S3A**). This domain contains several putative phosphorylation sites, 6 of which fit the minimal CDK phosphorylation motif ([S/T]-P) and 5 of which are conserved among OMA proteins from *Caenorhabditis* species (**Fig. S3A**). Moreover, OMA-1’s C-terminal region is sufficient to induce its degradation *in vivo* (Beer et al., 2019).

To test whether CYB-3’s PBP induces OMA degradation by targeting its C-terminus, we generated a transgene expressing OMA-2’s amino acids 208-393 fused to mNeonGreen, under a germline promoter (*Pmex-5*) and the *oma-2* 3’UTR (referred to as mNG::OMA-2^CT^) (**Fig. 4A**). We found that, similar to full-length OMA-2, OMA-2^CT^ degraded in early embryos at the onset of PNM, although with slower kinetics when compared to full length OMA-2 (**Fig. 4A& B; S3B& C**). These slower kinetics may either reflect contributions from other OMA domains to timely degradation, or higher expression levels of the C-terminal fusion construct compared to full length OMA-2. Nevertheless, mutating the CYB-3 PBP delayed OMA-2^CT^ degradation (**Fig. 4A& B; S3B&C**), confirming that OMA-2’s C-terminal region is required for its CYB-3–CDK-1’s dependent post-PNM degradation.

**Figure 4.**
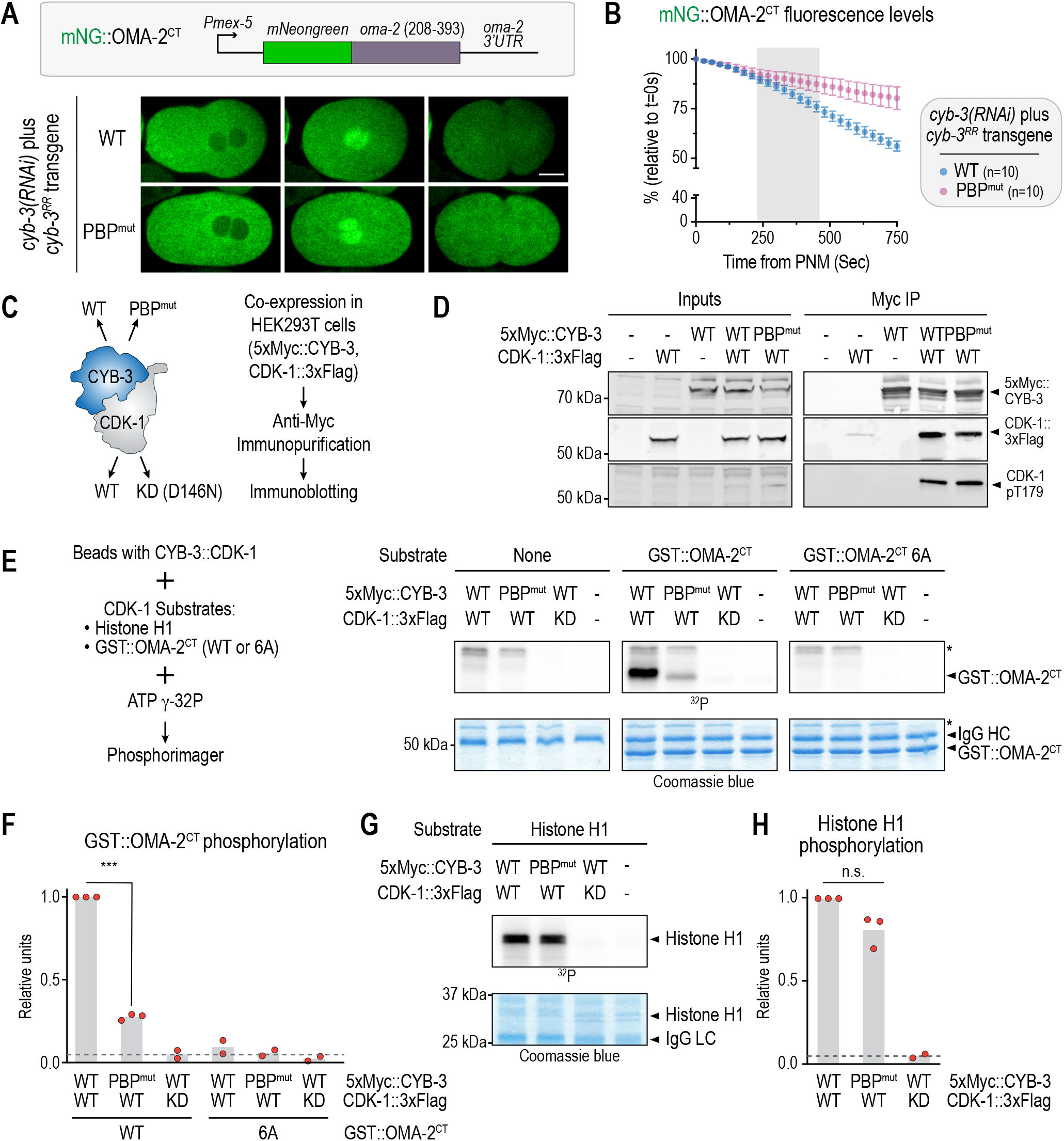
The CYB-3 PBP is critical for CDK-1 – dependent OMA hyperphosphorylation. **(A)** *(top)* Schematic illustrating an OMA-2 degradation reporter corresponding to the C-terminus of OMA-2 (amino acids 208-393) fused to mNeonGreen (mNG::OMA-2^CT^) and driven by a *mex-5* promoter (*Pmex-5*) and *oma-2* 3’UTR. *(bottom)* Stills from time-lapse sequences of embryos expressing mNG::OMA-2^CT^ and the corresponding *cyb-3* transgenes, undergoing early embryogenesis. Scale bar, 10 µm. **(B)** Quantification of mNG::OMA-2^CT^ fluorescence intensity levels for the conditions specified in *(A)*. Error bars correspond to 95% confidence intervals. *See also Fig. S3B&C*. **(C)** *(left)* Schematic illustrating the CYB-3–CDK-1 complex and the corresponding mutants tested in *in vitro* kinase assays. *(right)* Strategy used for co-expression of CYB-3 and CDK-1 in HEK293T cells and immunopurification of complexes. **(D)** Immunoblot of CYB-3–CDK-1 complexes expressed in HEK293T cells and immunopurified using a resin coupled to anti-Myc antibodies. CDK-1 pT179 is equivalent to human CDK1 pT161 and marks T-loop phosphorylated CDK-1. **(E)** *(left)* Strategy used for *in vitro* kinase assays. *(right) In vitro* kinase assays using CYB-3–CDK-1 complexes immunoprecipitated from HEK293T cell extracts, using GST::OMA-2^CT^ WT or 6A as substrates. IgG HC corresponds to the immunoglobulin heavy chain, which was present in the anti-Myc resin. Asterisks denote a band potentially corresponding to 5xMyc-tagged CYB-3 undergoing auto-phosphorylation. **(F)&(H)** Quantification of kinase activity for the specified conditions. Dashed lines represent baseline phosphorylation activity using kinase deficient CDK-1. **(G)** *In vitro* kinase assays using CYB-3–CDK-1 complexes immunoprecipitated from HEK293T cell extracts, using Histone H1 as a substrate. IgG LC corresponds to the immunoglobulin light chain, which was present in the anti-Myc resin. *n* represents the number of embryos analyzed. *** represents *P* < 0.001 from Welch’s t-tests; non-significant (n.s.) is *P* > 0.05.

Work from other systems has demonstrated that the PBP in cyclin B proteins plays a key role in targeting CDK substrates for hyperphosphorylation by working as a processivity mechanism that drives multi-site phosphorylation (Asfaha et al., 2022; Heinzle et al., 2025; Ng et al., 2025; Schunk et al., 2025). To test whether the CYB-3 PBP operates in a similar manner, we developed an *in vitro* kinase assay. To facilitate the production of active CYB-3–CDK-1 complexes, we co-expressed 5xMyc-tagged CYB-3 and 3xFlag-tagged CDK-1 in HEK293T cells, to then isolate the resulting complexes through immunopurification (**Fig. 4C**). Under these conditions, 5xMyc::CYB-3 co-immunopurified with CDK1::3xFlag (**Fig. 4D; S3D**). Importantly, in this heterologous system, CYB-3 – bound CDK-1 is phosphorylated at Thr179 (equivalent to pT161 in human CDK1) (**Fig. 4D**), a critical site for the activation of CDK-1 (Russo et al., 1996). Having validated the reconstitution of CYB-3–CDK-1 complexes in HEK293T cells, we next tested their ability to phosphorylate OMA *in vitro*. For this, we expressed and purified a fragment containing OMA-2’ C-terminus (amino acids 208-393) as a GST fusion (referred to as GST::OMA-2^CT^) (**Fig. S3E**). Notably, while CYB-3–CDK-1 was able to efficiently phosphorylate GST::OMA-2^CT^, mutating the PBP in CYB-3 reduced the ability of CDK-1 to target GST::OMA-2^CT^ for phosphorylation to ∼24% (**Fig. 4E&F**). Importantly, either mutating the 6 putative CDK sites in GST::OMA-2^CT^ to alanines or mutating CDK-1 to render it kinase deficient (KD, D146N; **Fig. 4C**) abolished this phosphorylation (**Fig. 4E&F**), indicating that the CYB-3–CDK-1 complex hyperphosphorylates OMA-2 in a PBP-dependent manner. On the other hand, both CYB-3 WT–CDK-1 and CYB-3 PBP^mut^–CDK-1 complexes were similarly efficient in their ability to phosphorylate Histone H1, a canonical CDK substrate (**Fig. 4G&H**), suggesting mutating the PBP does not substantially impair CYB-3’s ability to activate CDK-1. We conclude that the CYB-3 PBP is essential for CDK-1-dependent OMA hyperphosphorylation.

### Identification of a potential docking site in OMA for cyclin B3’s PBP

The PBP in cyclin B is proposed to dock onto pre-phosphorylated substrates to drive their hyperphosphorylation (Asfaha et al., 2022; Heinzle et al., 2025; Ng et al., 2025; Schunk et al., 2025). To gain further insight into the molecular mechanism of CYB-3 PBP-dependent OMA degradation, we asked whether any of the six putative CDK phosphorylation sites in OMA-2 — T227, T240, S291, S314, T327, and S335 — might function as a docking site for CYB-3’s PBP (**Fig. 5A**). The OMA-1 equivalents to two of these sites, T227 and T327, have been shown to be required for its degradation and are predicted targets of MBK-2 and GSK-3, respectively (Nishi and Lin, 2005) (**Fig. 5A**). To test the importance of putative CDK sites on OMA degradation, we took an unbiased approach by mutating each of the six putative CDK sites to alanines. To facilitate these assays while avoiding potential lethality effects associated with stabilization of OMA-2, we used our mNG::OMA-2^CT^ transgenic system as a reporter for OMA degradation. Strikingly, mutation of either T227 or T240 alone was sufficient to stabilize the protein to a degree comparable to mutating the CYB-3 PBP (**Fig. 5B&C**) and to stabilize it in late-stage embryos (**Fig. S4A**). On the other hand, mutating the remaining sites individually resulted in partial OMA-2 stability, with the non-conserved S335 site having the strongest effect (**Fig. 5B&C**) (note that mutation of S335 did not stabilize OMA-2^CT^ in late-stage embryos, **Fig. S4A**). While the reduced OMA-2 degradation caused by mutation of T227 is consistent with this site being a critical target of MBK-2, T240 has not been previously shown to be required for OMA degradation. Thus, we hypothesized that T240 is a potential CYB-3–CDK-1 complex phosphorylation and docking target. To test this, we returned to our kinase assays and we found that mutating T240 in GST::OMA-2^CT^ reduced its phosphorylation to ∼28% of wild-type (**Fig. 5D&E**), which is comparable to the reduction observed upon mutation of the CYB-3 PBP. On the other hand, mutating T227 had no effect on CYB-3–CDK-1 complex – dependent OMA-2 hyperphosphorylation (**Fig. S4B**). Combining CYB-3 PBP^mut^ with GST::OMA-2^CT^ T240A further reduced OMA-2 phosphorylation to ∼12% (**Fig. 5D&E**). These data are consistent with T240 being a primary OMA-2 site targeted by the CYB-3–CDK-1 complex to drive OMA hyperphosphorylation, although additional sites may also exist.

**Figure 5.**
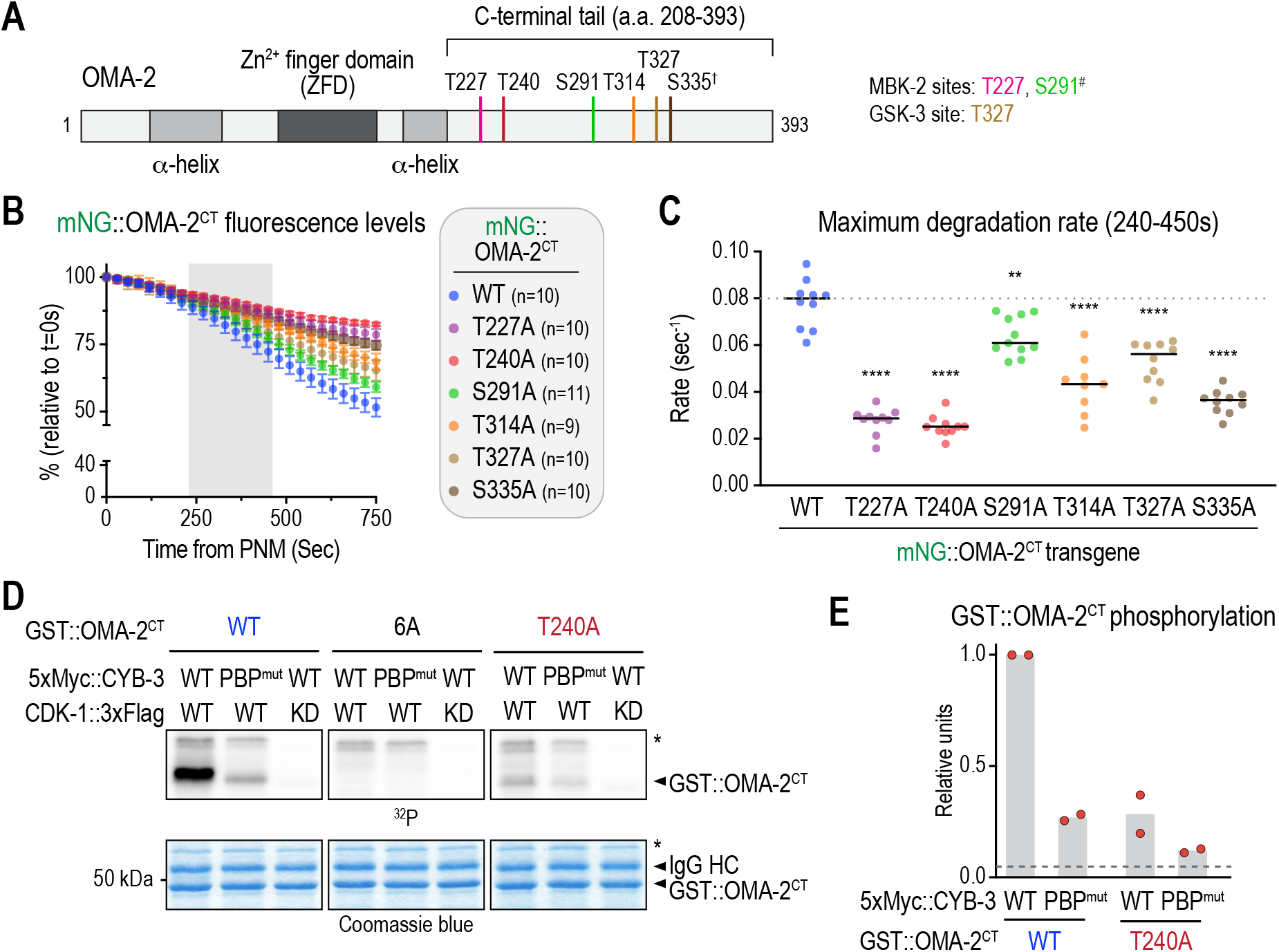
Identification of a potential OMA site for the docking of CYB-3’s PBP to drive its hyperphosphorylation. **(A)** Schematic illustrating the structure of OMA-2, as well as putative CDK phosphorylation sites in its C-terminal tail. (†) denotes a putative CDK site in OMA-1 that is not conserved as such in OMA-1, whereas (#) indicates a residue attributed to MBK-2 that does not contribute majorly to OMA degradation. **(B)** Quantification of mNG::OMA-2^CT^ fluorescence intensity levels for the specified mutants. Gray zone represents time of maximum OMA degradation, used for calculating degradation rates in *(C)*. Error bars are 95% confidence intervals. **(C)** Quantification of maximum degradation rate from slopes between 240-450 sec post-PNM from *(B)*. Dotted line indicates mean degradation rate for WT OMA-2^CT^. **(D)** *In vitro* kinase assays using CYB-3–CDK-1 complexes immunoprecipitated from HEK293T cell extracts and GST::OMA-2^CT^ (WT, 6A or T240A) as a substrate. IgH HC corresponds to the immunoglobulin heavy chain present in the anti-Myc resin. Asterisks denote a band potentially corresponding to 5xMyc-tagged CYB-3 undergoing auto-phosphorylation. **(E)** Quantification of the OMA-2 kinase activity for the conditions specified in *(D)*. Data for WT OMA-2^CT^ is the same as for Fig. 4E. Dashed line indicates baseline OMA-2^CT^ phosphorylation under CDK-1 kinase dead conditions (*see* Fig. 4F). *n* represents the number of embryos analyzed. **** represents *P* < 0.0001 from Mann-Whitney tests comparing each condition to the WT control; ** represents *P*< 0.01.

### Redundancy of phosphorylation sites ensures timely OMA degradation and normal embryonic development

Finally, we next sought to understand the relative contributions of individual phosphorylation sites to OMA degradation. For this, we introduced OMA-2 stabilizing mutants (T227A, T240A) to the endogenous locus, alone or in combination, using CRISPR-Cas9 (**Fig. 6A**). In order to prevent potential toxicity effects arising from these mutations, we generated a strain where OMA-2 was endogenously tagged with mNeonGreen and the auxin-induced degron (AID); these strains also expressed TIR1 in the germline (**Fig. 6A**). To suppress OMA-2 expression, auxin (20 µM 5-Ph-IAA) was maintained during the genome editing and clonal selection process and washed-out to allow OMA-2 levels to recover prior to phenotypic analysis (**Fig. 6A**) (note that endogenous expression of OMA-1 is sufficient to support oocyte development under these conditions; **Fig. S1A&B**). Consistent with our previous work using the OMA-2 degradation reporter, both T227A and T240A mutants significantly delayed OMA-2 degradation (**Fig. 6B-D**). However, mutation of T240, the CYB-3–CDK-1 phosphorylation site, exhibited a more pronounced effect than mutating T227 (**Fig. 6B-D**). In addition, mutating T227 and T240 in combination resulted in a small additive effect (**Fig. 6B-D**). Importantly, mutation of T240 resulted in ∼27% embryonic lethality, an effect that was enhanced to ∼46% when T227A and T240A were combined (**Fig. 6E**). Consistent with this embryonic lethality arising from OMA-2 stabilization, these effects were reversed by degrading OMA-2 through the addition of auxin (**Fig. 6E**). Surprisingly, mutation of T227 alone did not result in significant lethality (**Fig. 6E**), suggesting it did not stabilize OMA-2 to a level that was sufficient to produce detectable phenotypes. We conclude that multiple semi-redundant sites contribute to driving rapid OMA protein degradation to ensure normal embryonic development. Consistent with this, mutating the remaining residues that partially contribute to OMA-2^CT^ degradation (S291, T314, T327, and S335) stabilized OMA-2^CT^ to a greater extent than any of the single-site mutants and to an extent comparable to mutating T227 or T240 (**Fig. S4C**), supporting the notion that these residues act collectively as secondary phosphorylation sites.

**Figure 6.**
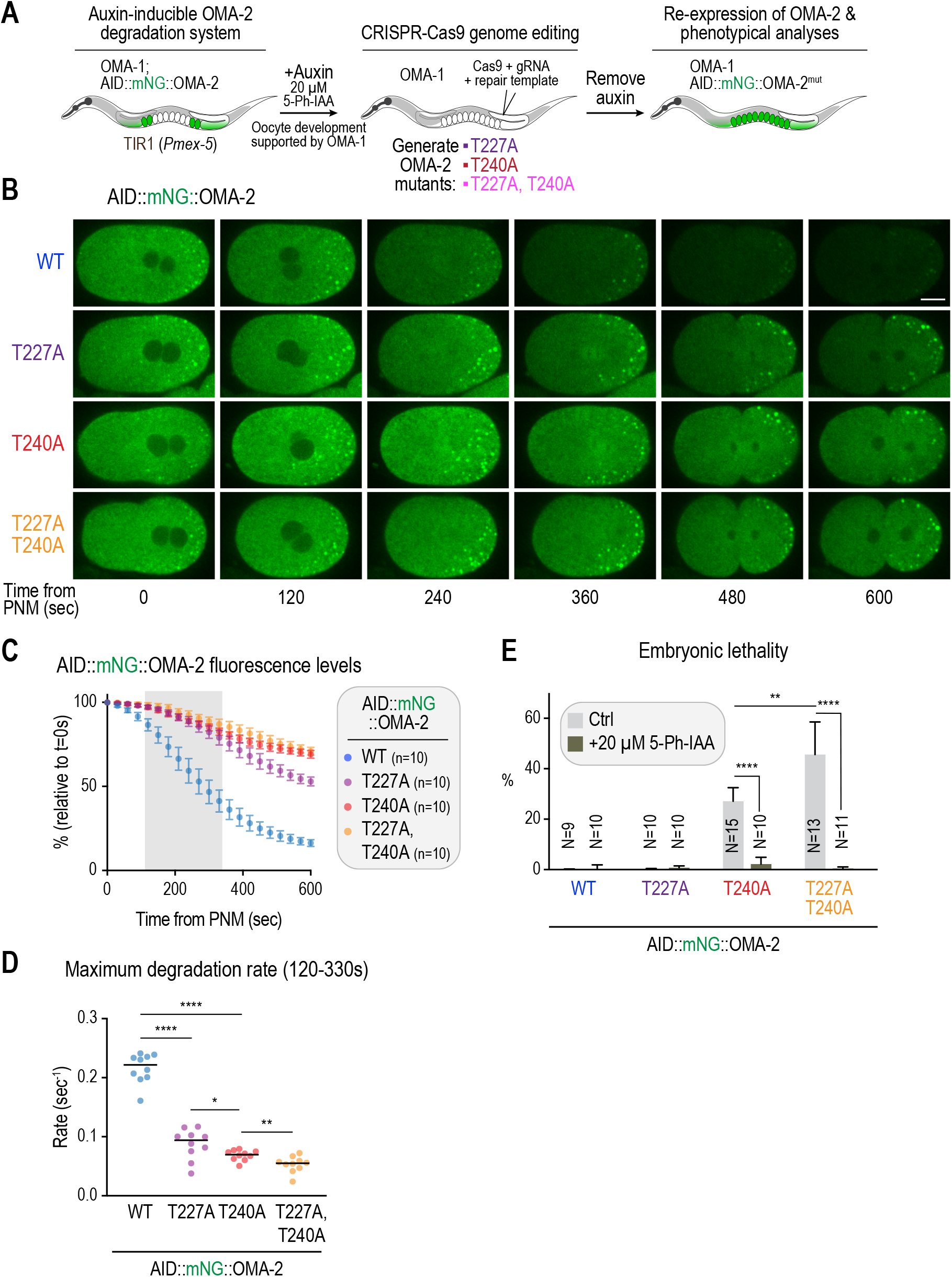
Two phosphorylation sites in OMA-2 are required for its degradation and for normal embryonic development. **(A)** Schematic illustrating the strategy utilized for the generation and analysis of OMA-2 endogenous mutants. **(B)** Stills from time-lapse sequences of embryos expressing AID::mNG::OMA-2, imaged under the specified conditions. Scale bar, 10 µm. **(C)** Quantification of AID::mNG::OMA-2 fluorescence intensity levels for the specified mutants. Gray zone represents time of maximum OMA degradation, used for calculating degradation rates in *(D)*. **(D)** Quantification of maximum degradation rate from slopes between 120-330 sec post-PNM from *(C)*. **(E)** Quantification of embryonic lethality values for the specified OMA-2 mutants, in the presence or absence of 20 µM auxin (5-Ph-IAA). *n* represents the number of embryos analyzed; *N* represents the number of adults whose progeny was scored. Error bars are 95% intervals. **** represents *P* < 0.0001 from Mann-Whitney tests; ** represents *P<* 0.01 and * represents *P<* 0.05.

## DISCUSSION

The results described here support a mechanism whereby multi-site phosphorylation of OMA, driven primarily by the phosphate binding pocket of CDK-1-bound CYB-3, ensures its rapid degradation during early embryogenesis (**Fig. 7**). Based on our data and previous work (Nishi and Lin, 2005), we propose a model where the unstructured C-terminus of OMA proteins is phosphorylated after meiosis II by MBK-2, followed by the CYB-3–CDK-1 complex. These primary sites, particularly the putative CDK-1 site (T240 in OMA-2), engage the phosphate binding pocket of CYB-3 to drive OMA phosphorylation at secondary sites, which create phospho-degrons that target OMA for degradation. This degradation serves to relieve mRNA repression and enable expression of early cell fate determinants (**Fig. 7**).

**Figure 7.**
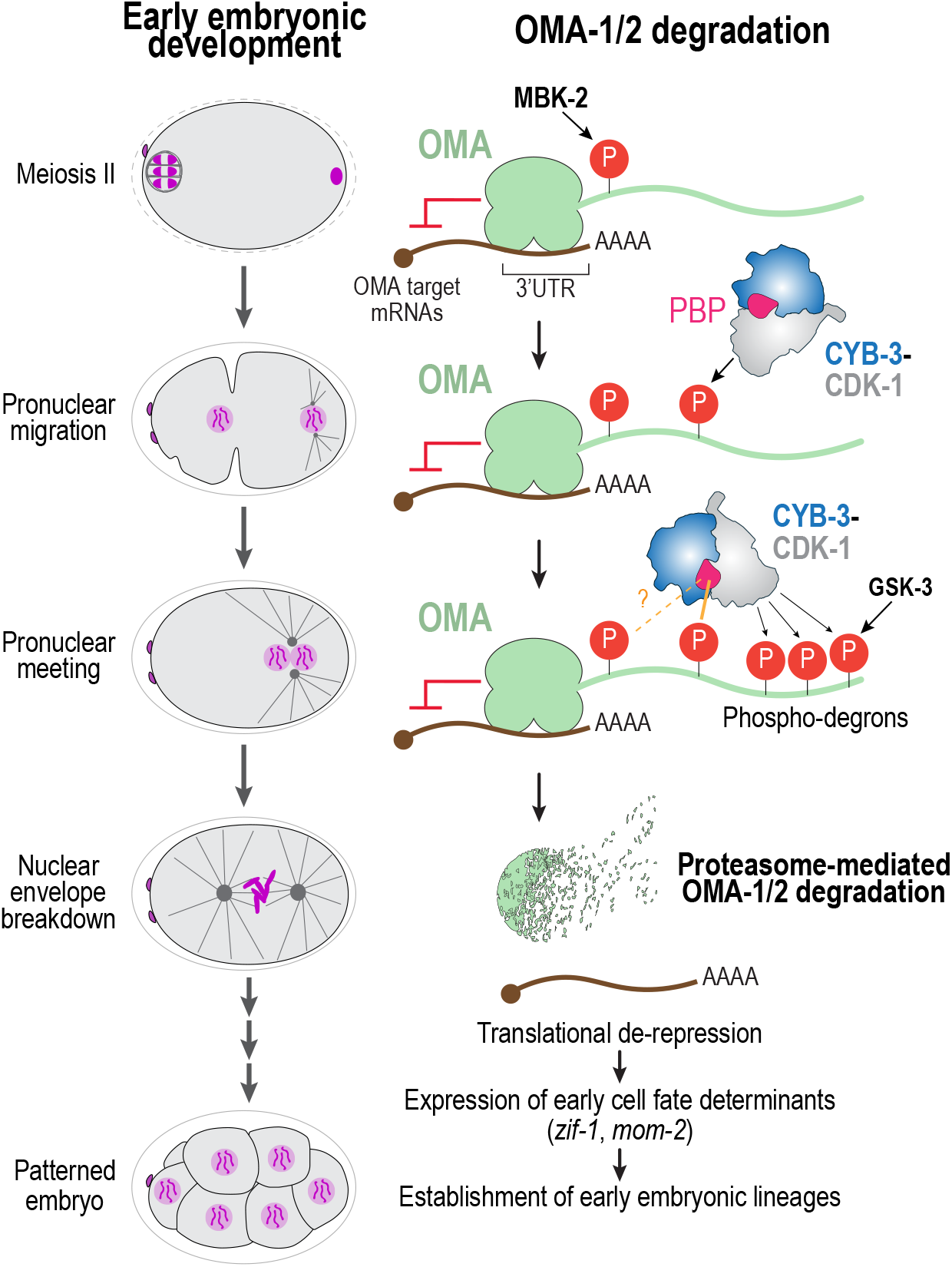
Proposed mechanism for OMA phosphorylation and degradation in early embryogenesis. Phosphorylation of OMA by MBK-2, which activates upon meiosis II completion, occurs first, followed by CYB-3–CDK-1. These initial phosphorylation events promote further, secondary OMA phosphorylation driven by the CYB-3 PBP. Secondary events create phospho-degrons that are recognized by ubiquitin ligases for the rapid degradation of OMA and the de-repression of maternal mRNAs.

### OMA hyperphosphorylation and degradation is mediated by the phosphate binding pocket of CYB-3

Our study provides a molecular mechanism for how the degradation of the oocyte protein OMA depends on the CYB-3–CDK-1 complex. During mitosis in *C. elegans* embryos, CDK-1 is activated by three cyclin B isoforms: CYB-1, CYB-2, and CYB-3 (van der Voet et al., 2009). Combined depletion of all three isoforms is required to recapitulate the CDK-1 loss-of-function phenotype and to preclude mitotic entry (Lara-Gonzalez et al., 2024; van der Voet et al., 2009). Despite their functional redundancy in driving mitosis, we found that CYB-1 and CYB-2 do not significantly contribute to OMA degradation, which is consistent with prior analyses (Shirayama et al., 2006). Instead, timely degradation of OMA specifically requires CYB-3 and its PBP. The molecular basis for this selectivity is not clear, as all three cyclin B isoforms possess predicted phosphate binding pockets (Yu et al., 2021). One possibility is that differences in pocket geometry or electrostatic environment determine substrate compatibility, such that phosphorylated OMA can be accommodated by the CYB-3 PBP, but not those of CYB-1 or CYB-2. In the case of vertebrate cyclin B3, its binding to phosphorylated targets also depends on the presence of a basic patch downstream of the phosphorylated residue (Schunk et al., 2025), although OMA proteins do not appear to contain such regions. Mass spectrometry analyses have identified several potential binding partners of the PBP of human cyclin B1 and yeast Clb2 (Heinzle et al., 2025; Ng et al., 2025), but no clear consensus binding motif has emerged from these studies. Interestingly, a prior genetic screen identified an I173F mutation in CDK-1 that stabilizes OMA (Shirayama et al., 2006), and its lethality effect was reported to be suppressed by a compensatory CYB-3 mutation, I116V (Ishidate et al., 2014). AlphaFold3 modelling predicts that these residues lie in the vicinity of the CYB-3 PBP (**Fig. S4D**), suggesting a shared binding surface between CDK-1 and CYB-3 involving specific amino acids. Further structural and biochemical work will be needed to define the sequence and structural requirements for PBP binding across cyclin B isoforms, and the sites characterized here may help inform those efforts.

Our finding that CYB-3–CDK-1 promotes OMA hyperphosphorylation and degradation via a phosphate binding pocket parallels work in vertebrate oocytes, where cyclin B3 promotes the PBP-dependent hyperphosphorylation of the APC/C inhibitor Emi2/XErp1 for its subsequent degradation to trigger meiosis I exit (Bouftas et al., 2022; Schunk et al., 2025). In this case, degradation also depends on Polo-like kinase 1 (Plk1), which docks onto phospho-Emi2/XErp1 to generate phospho-degrons that are recognized by the SCF^β-TRCP^ ubiquitin ligase complex (Wu and Kornbluth, 2008). By contrast, much less is known about the events in OMA degradation downstream of its phosphorylation and the phosphorylation sites identified so far do not appear to match the consensus sequence for a canonical phospho-degron (Zhang et al., 2025).

Interestingly, at least three ubiquitin ligase complexes, SCF^LIN-23^, SCF^FBXB-3^, and CRL2^ZYG-11^, have been proposed to promote OMA degradation (Du et al., 2015), which may indicate multiple parallel mechanisms for its degradation.

### Multi-site phosphorylation drives rapid OMA degradation in early embryogenesis

Prior work had established the contributions of two kinases to OMA degradation: MBK-2 and GSK-3 (Nishi and Lin, 2005; Shirayama et al., 2006). Of these, MBK-2 appeared to play the more significant role, as mutation of its key target site in OMA-1 (T239) severely delayed OMA degradation and resulted in 100% embryonic lethality (Lin, 2003; Nishi and Lin, 2005). However, we found that mutation of the equivalent site in OMA-2 (T227) resulted in partial stabilization with no significant embryonic lethality. We speculate that this difference is due to OMA-1 degrading at a slower rate than OMA-2, such that even partial stabilization of OMA-1 results in a more severe phenotype. Nonetheless, these data revealed that the contribution of MBK-2 alone to OMA degradation is partial and that OMA degradation can tolerate delays without severely compromising embryonic viability, as long as they do not disrupt gene expression timing and interfere with critical cell fate specification events.

In contrast, mutation of the CDK-1 phosphorylation site in OMA-2 identified in this study (T240) resulted in significantly greater stabilization and embryonic lethality. We interpret this as T240 being a major PBP docking site, the occupancy of which drives subsequent phosphorylation of secondary sites by CDK-1. Notably, combining mutation of OMA-2 at sites T227 and T240 resulted in an additive effect on OMA stability and increased embryonic lethality. In addition, combining mutations at sites in OMA-2 that individually caused modest stabilization resulted in greater stability than any single mutation alone. Together, these findings support a model in which the CYB-3–CDK-1 complex and MBK-2 cooperate to drive multi-site phosphorylation of OMA, which is required for its rapid degradation in early embryogenesis and for the timely expression of early cell fate determinants.

OMA proteins are members of the ZFP36 family of zinc finger proteins. Interestingly, Cth1, a member of this family, was recently shown to play a role in transcript regulation during zebrafish embryogenesis (Kushawah et al., 2024). Sequence analysis reveals putative CDK phosphorylation sites within Cth1’s unstructured C-terminal region, suggesting that Cth1 may be subject to a similar CDK1-dependent regulatory mechanism, although this remains to be explored.

### Separating CYB-3–CDK-1’s mitotic function from its role in OMA degradation

Our prior analysis of cyclin B contributions to *C. elegans* embryogenesis revealed that CYB-3 is essential for accelerating the pace of early mitoses, acting as a more potent activator of CDK-1 than CYB-1 (Lara-Gonzalez et al., 2024). Accordingly, depletion of CYB-3 caused delays in chromosome alignment, spindle assembly, and chromosome segregation, along with a higher rate of mitotic chromosome segregation errors, which resulted in penetrant embryonic lethality (Lara-Gonzalez et al., 2024). By contrast, we found that when we mutate the phosphate binding pocket in CYB-3, the CYB-3–CDK-1 complex can still support mitosis and efficiently phosphorylate canonical substrates such as Histone H1. Even though CYB-3 PBP^mut^ resulted in mild mitotic delays, these alone are not predicted to result in mitotic errors or developmental defects (Houston et al., 2023). Nevertheless, CYB-3 PBP^mut^ embryos were fully embryonic lethal. Therefore, this lethality likely arose from developmental defects downstream of OMA stabilization, such as delayed expression of mRNAs such as *zif-1* and *mom-2*. Thus, our data support the notion that the CYB-3–CDK-1 complex coordinates two essential events for early embryogenesis: rapid embryonic mitoses and early embryo mRNA translation. We propose that post-meiotic activation of CDK-1 ensures that cell division is coupled to early gene expression for the correct specification of blastomeres during development.

Taken together, our findings establish CYB-3–CDK-1-mediated hyperphosphorylation as a key step in targeting OMA for degradation and initiating translation in the early embryo, revealing a molecular link between the cell cycle machinery and the developmental gene expression program. This work has the potential to reshape our view of early embryogenesis, which may impact our understanding and treatment of developmental and congenital disorders.

## MATERIALS AND METHODS

### *C. elegans* strains growth and maintenance

The strains used in this study are listed in **Table S1**. *C. elegans* strains were maintained at 20°C in nematode growth media (NGM) plates treated with 50 μg/ml of Streptomycin and seeded with the OP50-1 strain of *Escherichia coli*.

### Single-copy insertion transgenes

When generating transgenic lines, we used Mos1 Single-Copy Insertion (MosSCI) method (Frokjaer-Jensen et al., 2008). For this, transgenes cloned into either pcFJ352 or pcFJ151 were injected onto strains EG67011 (chromosome I) or EG6699 (chromosome II) respectively, to create single copy insertions (Frokjaer-Jensen et al., 2008). Integrants were selected by the ability of the transgene to rescue the *unc* phenotype of the parental strain and were confirmed by PCR.

### Editing genes at the endogenous locus by CRISPR-Cas9

When generating endogenous tags and mutations, we used CRISPR-Cas9-mediated genome editing methods described before (Hattersley et al., 2018). Briefly, a ribonucleic protein (RNP) mix containing purified Cas9 (Macrolab, UC Berkeley), crRNA (**Table S2**), and tracrRNA (IDT) was injected into the gonad of gravid adult animals, along with repair templates encoding for fluorescent tag or discreet mutations. An RNP mix targeting the *dpy-10* gene was simultaneously injected as a co-CRISPR strategy. Clones were selected for by the presence of the dumpy and roller phenotypes. Mutations and insertions were confirmed via PCR followed by Sanger DNA sequencing and the *dpy-10* mutation was removed by crossing back to parental strain.

For generating endogenous *oma-2* T227A and T240A mutations, adults were transferred to NGM plates containing 20 µM 5-Ph-IAA (MedChemExpress #HY-134653) after injection and during clonal selection. To re-express OMA-2, L4-stage worms were transferred to regular NGM plates and analyzed one generation later.

### RNA interference (RNAi)

For double stranded RNA (dsRNA) generation, single stranded RNAs (ssRNAs) were first synthesized *in vitro* using PCR templates containing T3 and T7 RNA polymerase sites (Hattersley et al., 2018). The resulting ssRNAs were then hybridized, aliquoted and stored at - 80°C. The oligonucleotides used for creating PCR templates are listed in **Table S3**. For RNAi, dsRNAs were injected at a final concentration of 0.8-1 mg/ml into L4 larvae and allowed 36-48 h at 20°C to recover before imaging.

For embryonic lethality assays, injected worms were recovered for 24 hours and were subsequently seeded on individual 35-mm plates and allowed to lay progeny for further 24 hours before being removed. The next day, the number of living larvae and dead embryos were recorded.

### Live imaging microscopy

*C. elegans* embryos were dissected from gravid adults in M9 media, mounted onto 2% agarose pads, covered with a #1.5 glass coverslip and imaged at 20-21°C. Time-lapse imaging was performed using a spinning disk microscope run by a NIS-Elements AR software (Nikon), and coupled with a CSU-X1 spinning disk unit (Yokogawa) mounted on an inverted microscope (Ti2E, Nikon), 40x Plan Fluorite oil objective (1.3NA) with a 1.5x intermediate magnification module, and outfitted with an Ixon 888 electron multiplication back-thinned charged-coupled device Life camera (Andor Technology). 5 or 7 layer stacks at 5 µm steps were taken at 15, 20 or 30 sec intervals, with a 488 nm excitation laser and a 455/50M, 25MM emission filter for green, or a 561 nm laser and a 605/52M, 25MM emission filter for red.

### Imaging analysis

Imaging analysis was done on ImageJ, using maximum intensity projections. Pronuclear meeting (PNM) was defined as the first frame in which pronuclei were in contact with one another in the one-cell embryo. Nuclear envelope breakdown (NEBD) was defined as the frame where free histone signal in the nucleus equilibrated with the cytoplasm. In strains without histone markers, NEBD was defined as the frame in which cytoplasmic markers entered the nucleus. Anaphase onset was scored as the first frame with visible separation of sister chromatids.

To quantify endogenously tagged OMA protein and OMA reporter abundance, an area was drawn surrounding the whole embryo, and the integrated density of a maximum intensity projection of the whole embryo was measured. The integrated density of background was subtracted by copying the same area to a region without embryos. Percentage fluorescence was calculated by dividing corrected intensity values by the initial intensity at PNM and multiplying by 100%.

To quantify *mom-2* and *zif-1* translational reporter abundance, an area was drawn surrounding the P2 or ABal nucleus, respectively. The integrated density of the nucleus was measured for both the reporter and an mCherry-Histone marker. The background was subtracted by copying the same area to a region within the embryo without nuclei for both reporter and histone intensity. Reporter intensity was then divided by histone intensity for normalization. For *mom-2* reporter, analysis was done 1 frame before NEBD of the P2 nucleus. For the *zif-1* reporter, analysis was done on the ABal nucleus 6 minutes after the EMS cell anaphase onset.

### Immunoprecipitation of mNeonGreen-tagged CYB-3 from *C. elegans* extracts

*C. elegans* strains expressing mNeonGreen-tagged CYB-3 (WT, HP^mut^, PBP^mut^ or CB^mut^) were seeded onto 10 mm plates and grown for 4 days at 20°C when the majority of the population corresponded to gravid adults. Worms were washed off using M9 buffer (22 mM KH_2_PO_4_, 42 mM Na_2_HPO_4_, 86 mM NaCl, and 1 mM MgSO_4_•7H2O), collected by centrifugation at 600g for 3 min and transferred to 15 mL conical tubes. After washing once more, worm pellets were resuspended in 1.0 mL of lysis buffer (50 mM HEPES pH 7.4, 1 mM EDTA, 1 mM MgCl_2_, 100 mM KCl, 10% glycerol, 0.1% Triton X-100, 1 mM DTT, EDTA-free protease inhibitor cocktail (Sigma #11697498001)). Worms were lysed using a tip sonicator (Fisher Scientific) and cleared by centrifugation at 20,000 g for 30 min at 4°C. Cleared lysates were incubated with mNeonGreen-nAb Agarose Beads (Allele Biotech #ABP-nAb-MNGA100), overnight at 4°C with rotation. Beads were washed 4 times with Lysis buffer and eluted by resuspension in 2X sample buffer (116.7 mM Tris-HCl pH 6.8, 3.3% SDS, 200 mM DTT, 10% glycerol, bromophenol blue).

### Preparation of *C. elegans* extracts for immunoblotting

30 to 60 worms per condition were collected in M9 media (22 mM KH_2_PO_4_, 42 mM Na_2_HPO_4_, 86 mM NaCl, and 1 mM MgSO_4_•7H_2_O) containing 0.1% Tween 20. Samples were spun down and washed 4 times. After last wash, supernatant was removed until 60 μL remained. An additional 20 μL were removed and replaced with 20 μL of 6x SDS sample buffer (116.7 mM Tris-HCl pH 6.8, 3.3% SDS, 200 mM DTT, 10% glycerol, bromophenol blue). Samples were sonicated in a water bath sonicator at 70°C for 10 minutes and heated for 5 minutes at 100°C using a heat block. Sonication and boiling were repeated twice.

### Immunoblotting

For immunoblotting of worm lysates, the equivalent of 10 worms was loaded onto 4-20% Bis-Tris Gels (GenScript, #M00657). Proteins were transferred to a nitrocellulose membrane using the Trans-Blot Turbo Transfer System (Bio-Rad, #1704158), Membranes were blocked with intercept blocking buffer (LicorBio #927-60001) and incubated overnight with primary antibody diluted in intercept blocking buffer. Membranes were then washed with 0.1% TBS-T (20 mM Tris-HCl, 150 mM NaCl, 0.1% Tween 20) before a two hour incubation with secondary antibody in intercept blocking buffer, at room temperature. Membranes were washed with TBS-T and developed in a LI-COR imaging system.

Antibodies used for immunoblotting were: mouse anti-myc (clone 9E10, Sigma-Aldrich #M4439;), mouse anti-Flag (clone M2, Sigma-Aldrich #F3165), rabbit anti-CDK-1 PSTAIRE (Millipore #6923), rabbit anti-Cdk1 pT161 (equivalent to *C. elegans* pT179; Cell Signaling #9114S), mouse anti-α-tubulin (clone DM1A, Millipore #T6199), rabbit anti-CYB-3(Lara-Gonzalez et al., 2024), 800CW goat anti-rabbit IgG (LI-COR #926-32211), and Alexa Fluor 680 goat anti-mouse IgG light chain specific (Jackson Immunoresearch #115-625-174).

### Protein purification

The vector used for generating glutathione-S-transferase (GST) fusion proteins was pGEX-LB, a derivative of pGEX-4T-1. In pGEX-LB, an encoded Pro residue is replaced with a Gly-Gly-Gly-Gly-Gly-Ser-Gly linker sequence to promote the independent functioning of the GST and fusion moieties (Bardwell et al., 2022).

GST fusion proteins were expressed in bacteria, purified by affinity chromatography using glutathione-Sepharose, eluted from the matrix with reduced glutathione, dialyzed to remove the glutathione, quantified by Bradford assay, and flash frozen in aliquots at 80°C, essentially as described elsewhere (Gordon et al., 2013). The substrate dialysis buffer was 20 mM Tris-HCl (pH 7.5), 0.1 mM EDTA, 1 mM dithiothreitol, 5% glycerol.

### Expression and purification of proteins from HEK293T cells

cDNAs encoding for CYB-3 and CDK-1 were cloned onto pcDNA3.1-based vectors encoding for 5xMyc or 3xFlag tags, respectively (**Table S4**). HEK293T cells were grown in 2×10 cm dishes per condition and transfected at 60% confluency with 10 µg of DNA constructs per plate using Lipofectamine 3000 Transfection Reagent and serum-free DMEM medium, according to product guidelines (Thermo Fisher Scientific #L3000001). Transfected cells were incubated for 48 h at 37°C and 5% CO_2_. To collect, cells were washed with Dulbecco’s phosphate buffer saline (ThermoFisher Scientific #14190-144) and resuspended in 1 mL of lysis buffer (0.1% Triton X-100, 10 mM Tris-HCl pH 7.4, 100 mM NaCl, 1 mM EDTA, 1 mM DTT, 20 mM β-glycerol phosphate, 10 mM NaF, 1 mM DTT, 0.2 mM PMSF and EDTA-free protease inhibitor cocktail; Sigma #11697498001). Cell suspensions were rotated for 30 minutes at 4°C before centrifugation at 13,000 rpm for 30 min at 4°C. For input, 50 µL of supernatant was taken and mixed with 50 µL of 4X SDS sample buffer and boiled for 10 minutes. The remaining supernatant was incubated with anti-Myc magnetic beads (Thermo Fisher Scientific #88842) equilibrated with lysis buffer, overnight at 4°C on an end-over-end rotator. After incubation, beads were washed 3 times with lysis buffer, resuspended in 1 mL of lysis buffer and used for kinase assays within 24 hr. For immunoblotting, 20 µL of bead sample was combined with 20 µL of 4X SDS-sample buffer and boiled for 10 minutes.

### Protein kinase reactions

Protein kinase reactions (20 µL) contained kinase assay buffer (50 mM Tris-HCl (pH 7.5), 10 mM MgCl_2_, 0.1 mM EGTA, 2 mM dithiothreitol, 0.01% Brij 35), purified substrate (1 µM Histone H1 (Sigma-Aldrich, cat #14-155), 5 µM GST::OMA-2^CT^ (or an equivalent volume of substrate dialysis buffer for “no substrate” points), 50 µM ATP, 1 µCi of [γ-^32^P]ATP, and ∼20 µL bed volume of bead-bound immunopurified CYB-3–CDK-1 (or control) complex that had been washed briefly with kinase assay buffer. Reactions were for incubated 30 min at 30°C, then stopped with SDS sample buffer, separated by 10% or 12% SDS-PAGE, and visualized and quantified on an Amersham Typhoon TRIO+ Imager using phosphor imaging mode.

## Statistical analyses

Statistical analysis was performed using Prism (Graphpad). Asterisks in figures denote statistical significance as estimated by Mann–Whitney tests. For figures 4F and 4H, a Whelch’s t-test was used. *, *P* < 0.05; **, *P* < 0.01; ***, *P* < 0.001; ****, *P* < 0.0001.

## ACKNOWLEDGEMENTS

We thank Sean Ryder for the GFP::OMA-1 strain, and Rebecca Green and Midori Ohta for their helpful comments on our manuscript. Some strains were provided by the CGC, which is funded by NIH Office of Research Infrastructure Programs (P40 OD010440). P.L.-G. is funded by an NIH R35 grant (GM150786). D.B. is funded by a Graduate Assistance in Areas of National Need (GAANN) fellowship.

## SUPPLEMENTARY FIGURE LEGENDS

**Figure S1.**
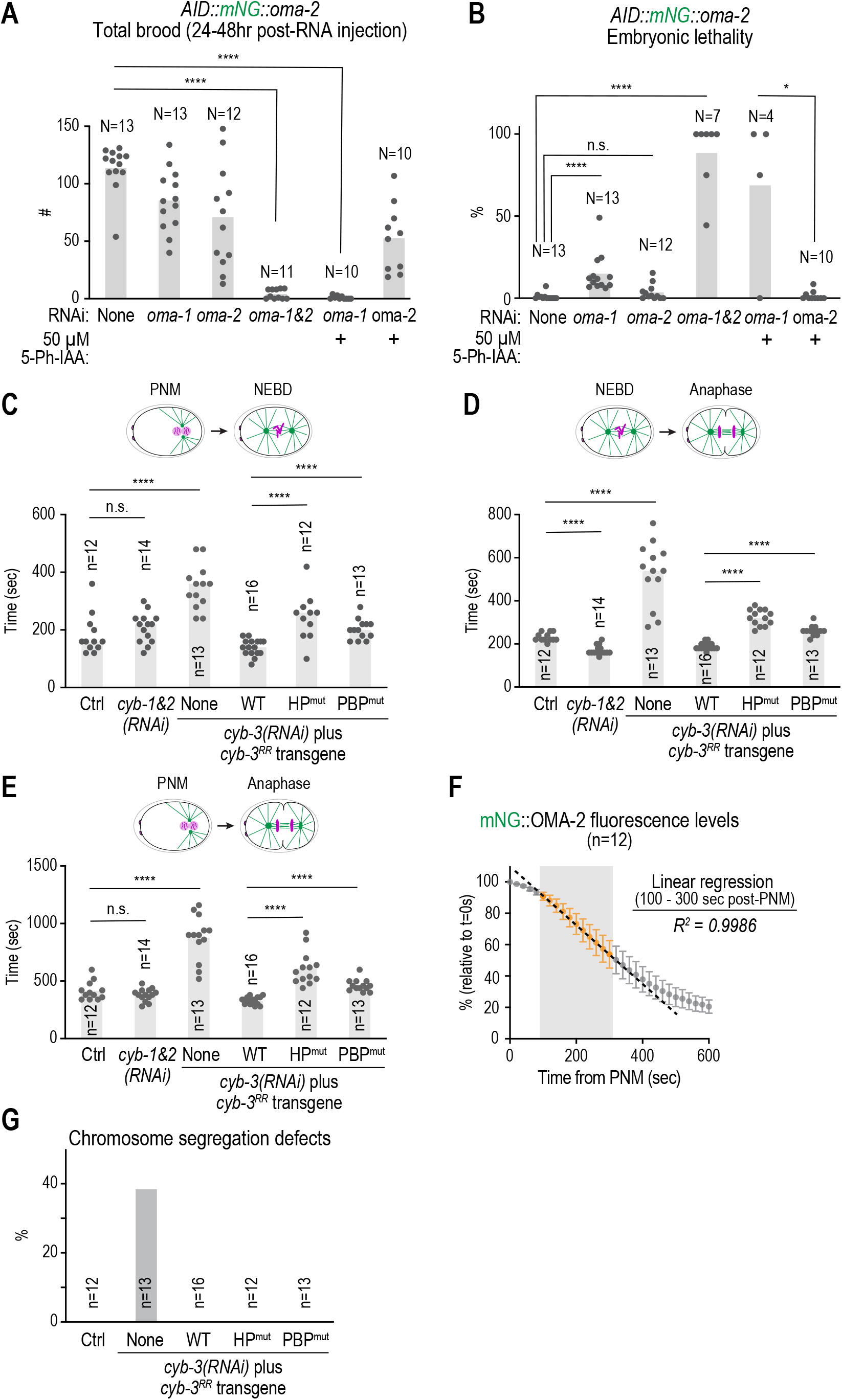
Functionality of mNeonGreen-tagged OMA-2 and the effect of CYB-3 mutants in cell cycle progression. *Related to* Fig. 1 and 2. (A)&(B) Quantification of total number of progeny laid *(A)* or embryonic lethality *(B)* in a strain expressing endogenously-tagged AID::mNG::OMA-2, under the specified conditions. Note that co-depletion of OMA-1 and OMA-2, either through RNAi or through the AID system, results in penetrant sterility, whereas depletion of OMA-1 in this background does not, indicating that AID::mNG::OMA-2 is functional. For panel *B*, adults with no progeny were excluded from the embryonic lethality analyses. **(C),(D)&(E)** Quantification of the interval from pronuclear meeting (PNM) to nuclear envelope breakdown (NEBD) *(C)*, from NEBD to Anaphase onset *(D)* and from PNM to Anaphase onset *(E)* under the specified conditions. Data for *(D)* are the same as for Fig. 2G. **(F)** Linear regression analysis for mNG::OMA-2 degradation for the data points labelled in orange, which fall within the gray shaded area (100-300 sec post-PNM). Note that the degradation rates within this zone fit a linear curve with high confidence (*R^2^ = 0.9986*) and were used to calculate maximum degradation rates (Fig. 1F, 2E, 6C). Data points are the same as for Fig. 1F. **(G)** Quantification of chromosome segregation defects under the specified conditions. *n* represents the number of embryos analyzed; *N* represents the number of adults whose progeny was scored. Error bars are 95% intervals. **** represents *P* < 0.0001 from Mann-Whitney tests; non-significant (n.s.) is *P* > 0.05.

**Figure S2.**
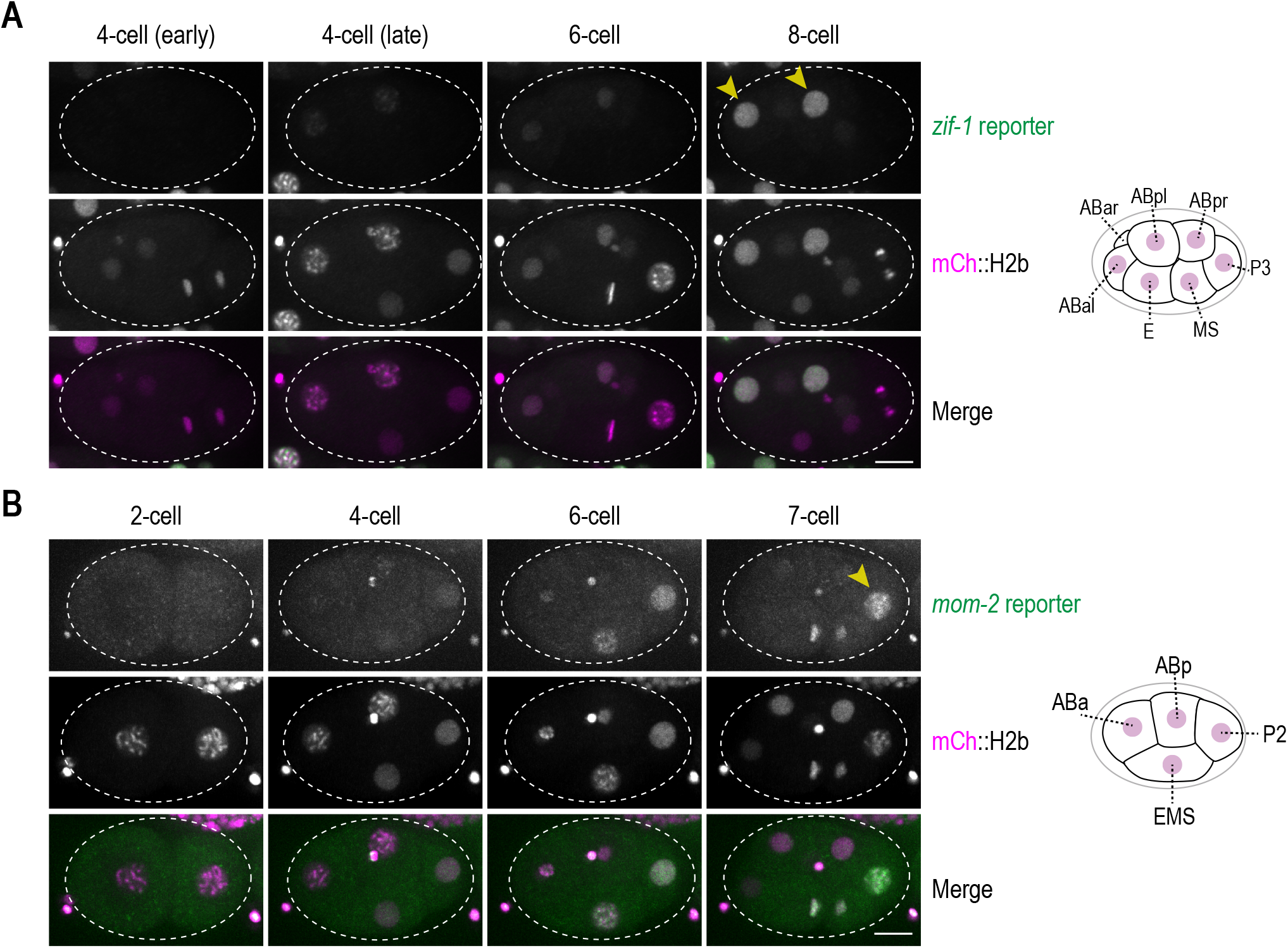
**Expression of translational reporters in early embryogenesis. *Related to*** Fig. 3**. (A)&(B)** Time-lapse sequences of embryos expressing mCherry-tagged Histone H2B, as well as *zif-1 (A)* or *mom-2 (B)* translational reporters. Note that the expression of both reporters is low at the 2-cell stage, but their translation increases significantly during the 4-8 cell stage. Arrowheads indicates the ABal, ABpl *(A)* or P4 *(B)* precursors, where expression for the respective reporters is maximum. Cartoons illustrating embryonic lineages are shown for reference. Scale bar, 10 µm.

**Figure S3.**
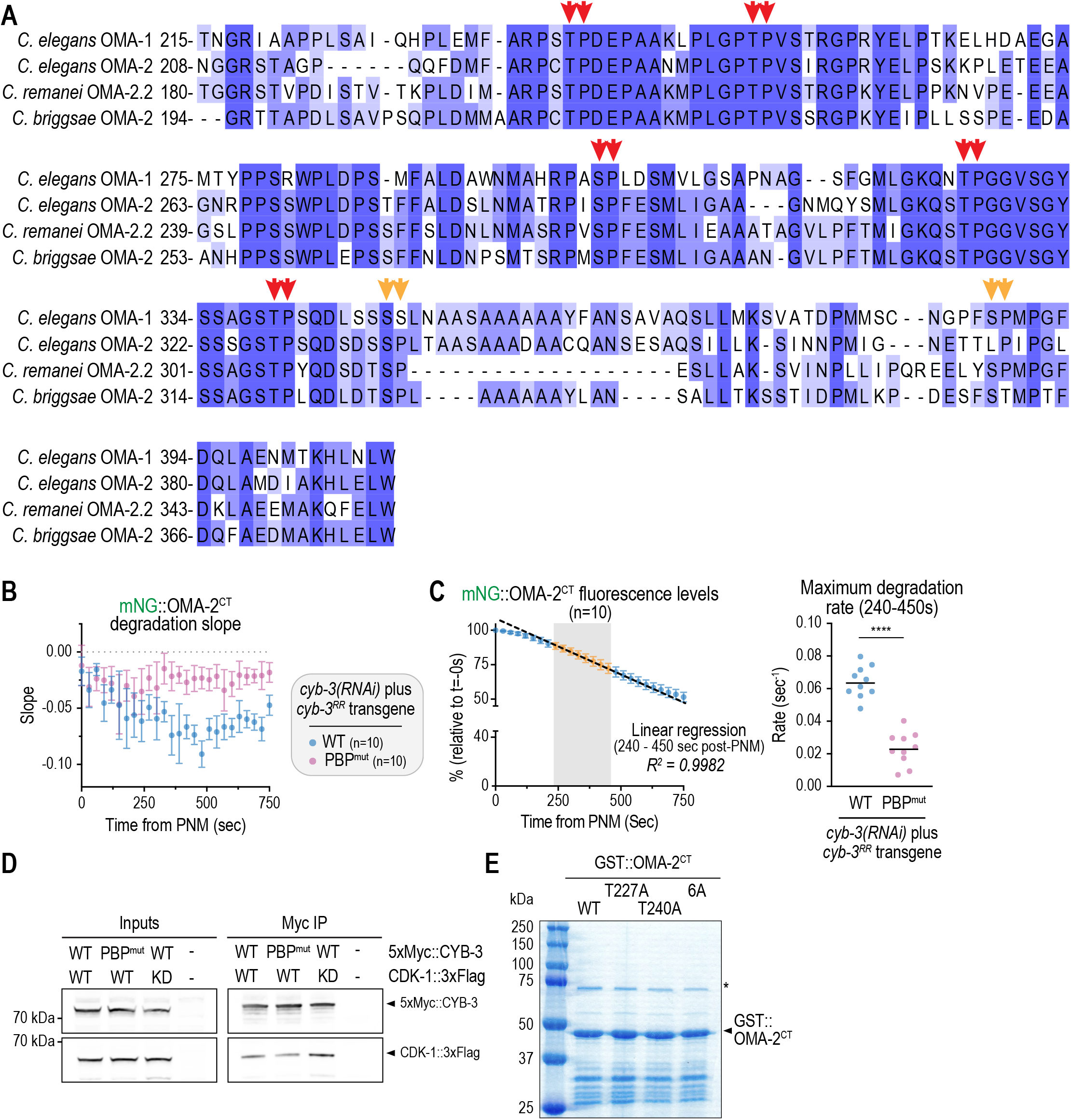
**Analysis of OMA’s C-terminus and generation of components for *in vitro*kinase assays. *Related to*** Fig. 4**. (A)** Alignment of C-terminal regions of *C. elegans* OMA-1 and OMA-2, as well as equivalent regions for OMA orthologues from *C. remanei* and *C. briggsae*. Double red arrowheads denote conserved CDK sites (SP or TP), whereas double yellow arrowheads show partially conserved sites. **(B)** Second derivative analyses for the data in Fig. 4B. **(C)** *(left)* Linear regression analysis for mNG::OMA-2^CT^ degradation for the data points labelled in orange, which fall within the gray shaded area (100-300 sec post-PNM). Note that the degradation rates within this zone fit a linear curve with high confidence (*R^2^ = 0.9982*) and were used to calculate maximum degradation rates (Fig. 5C*, S4C*). Data points are the same as for Fig. 5B. *(right)* Quantification of maximum degradation rate from slopes between 240-450 sec post-PNM from Fig. 4B. **(D)** Expression and immunopurification of CYB-3–CDK-1 complexes from HEK293T cells. **(E)** SDS-Polyacrylamide gel stained with Coomassie blue showing purified GST::OMA-2^CT^ either WT, 6A, T227A or T240A. Asterisk denotes a band corresponding to bovine serum albumin (BSA) that was used to block glutathione Sepharose beads used to purify GST-tagged proteins from bacteria and that is partially eluted.

**Figure S4.**
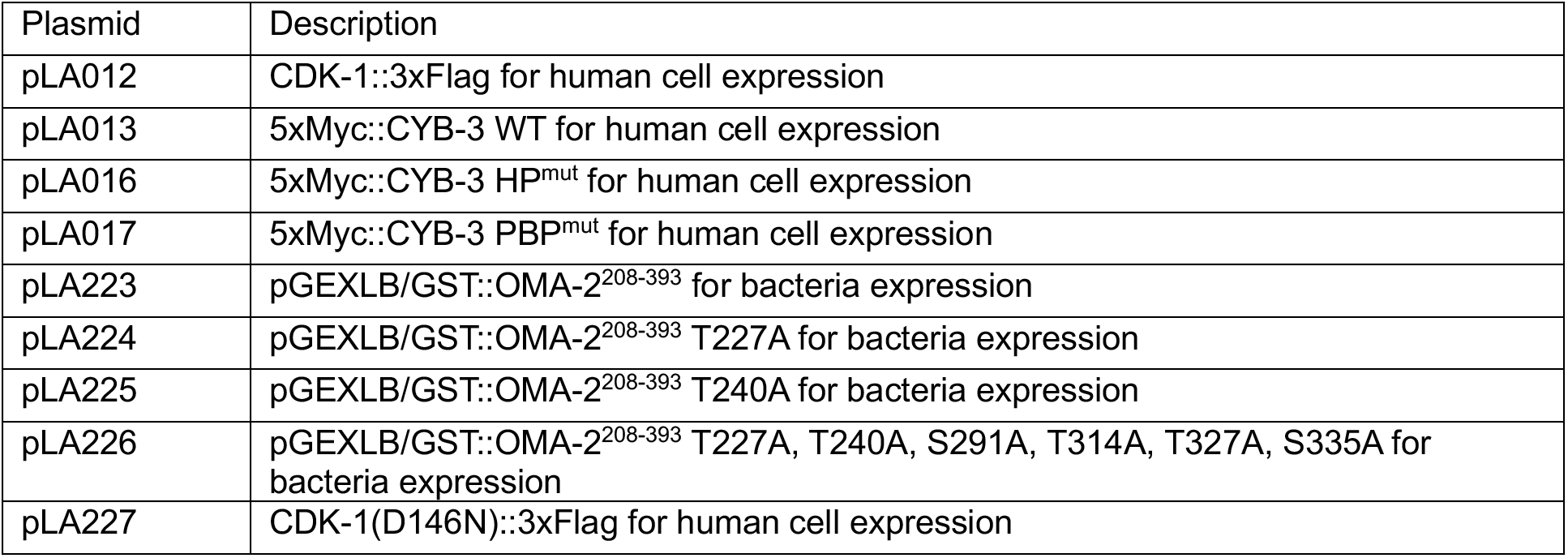
Analysis of OMA-2 phosphorylation mutants. *Related to* Fig. 5 and 6. **(A)** Whole adult images of strains expressing mNG::OMA-2^CT^ transgenes. Boxed area illustrates late-stage embryos within each adult. Whereas the WT transgene shows little expression in late-stage embryos, mutants exhibit mNG::OMA-2^CT^ with varying intensities in late embryos, indicating reduced degradation. Scale bar, 200 µm. **(B)** *In vitro* kinase assays using CYB-3–CDK-1 complexes immunoprecipitated from HEK293T cell extracts and GST::OMA-2^CT^ (WT or T227A) as a substrate. IgH HC corresponds to the immunoglobulin heavy chain present in the anti-Myc resin. Asterisks denote a band potentially corresponding to 5xMyc-tagged CYB-3 undergoing auto-phosphorylation. **(C)** Quantification of maximum degradation rate from slopes between 240-450 sec post-PNM for the specified mNG::OMA-2^CT^ mutants. Data for WT, S291A, T314A, T327A and S335A is the same as for Fig. 5C. **** represents *P* < 0.0001 from Mann-Whitney tests; *** is *P* > 0.001. **(D)** AlphaFold3 model of the CYB-3–CDK-1 complex displaying two key PBP residues in CYB-3: Arg250 and Arg267 (magenta). Mutation of CDK-1’s Ile173 (green) to phenylalanine was previouslyfound to result in stabilized OMA-1 (Shirayama et al., 2006), while mutation of CYB-3’s Ile116 (orange) to valine was reported to suppress this effect (Ishidate et al., 2014). Note that all these residues are predicted to be in the vicinity of the CYB-3 PBP and thus may form part of the same interface.

## SUPPLEMENTARY TABLES

**Table S1:**
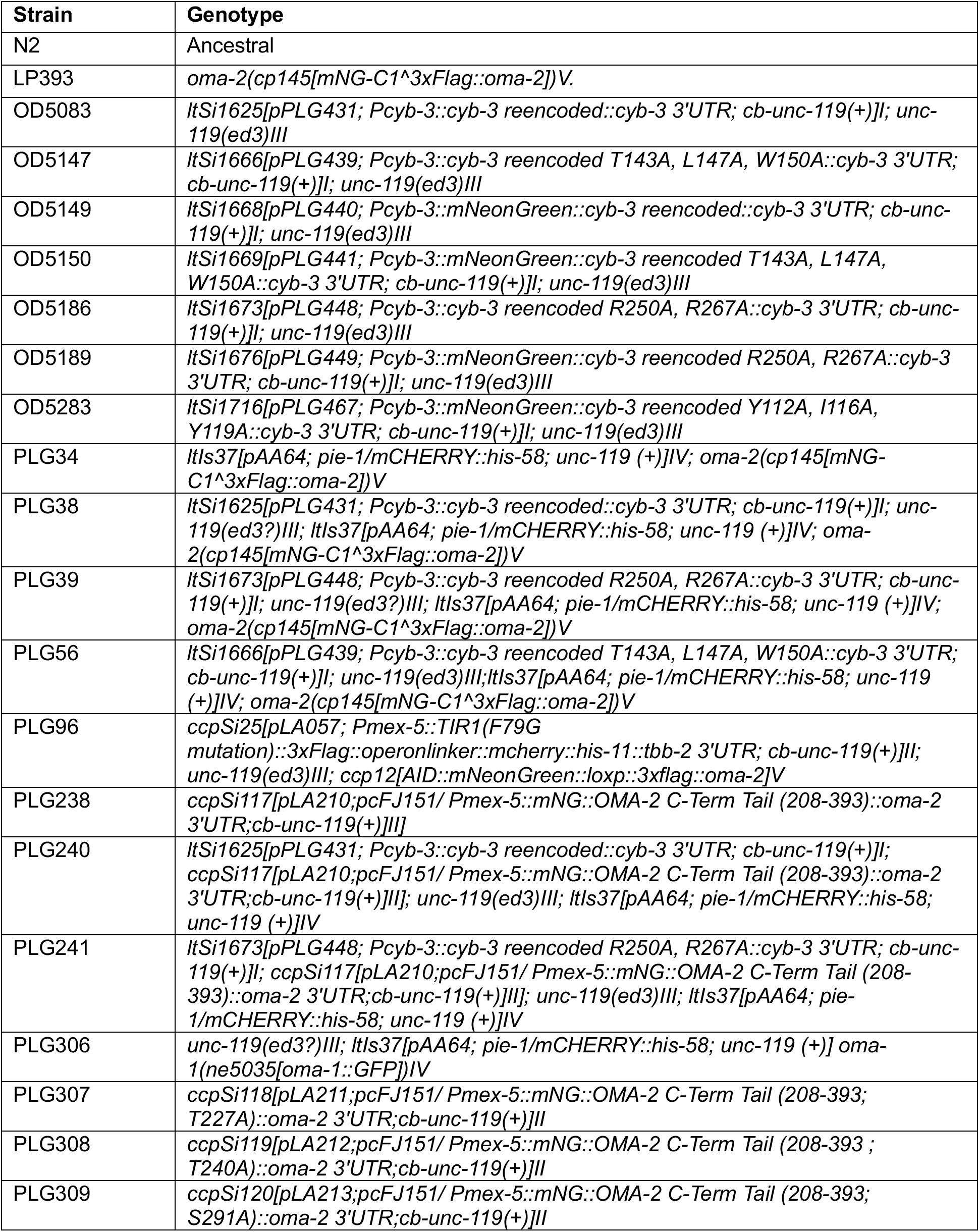
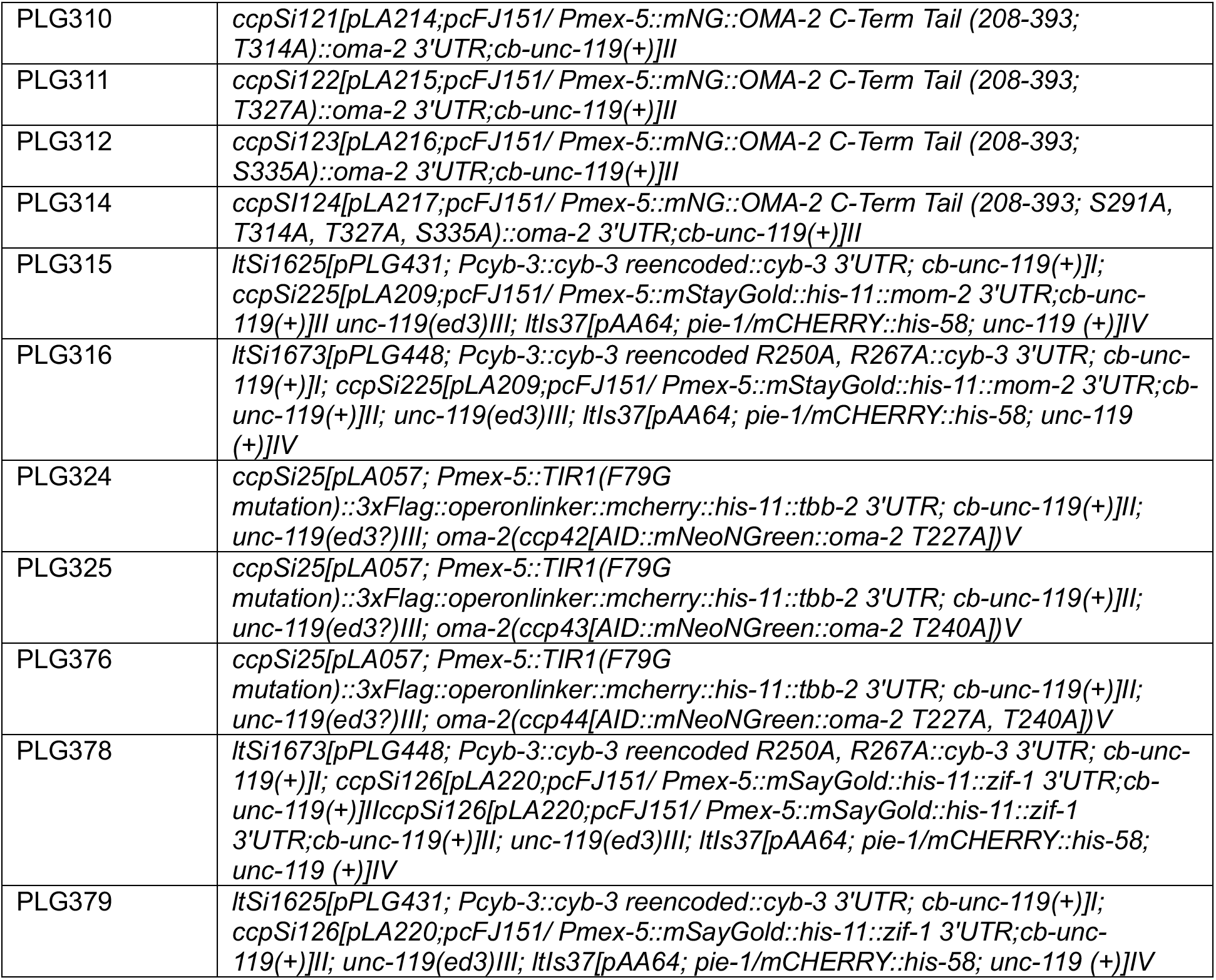
List of *C. elegans* strains used in this study.

**Table S2:**
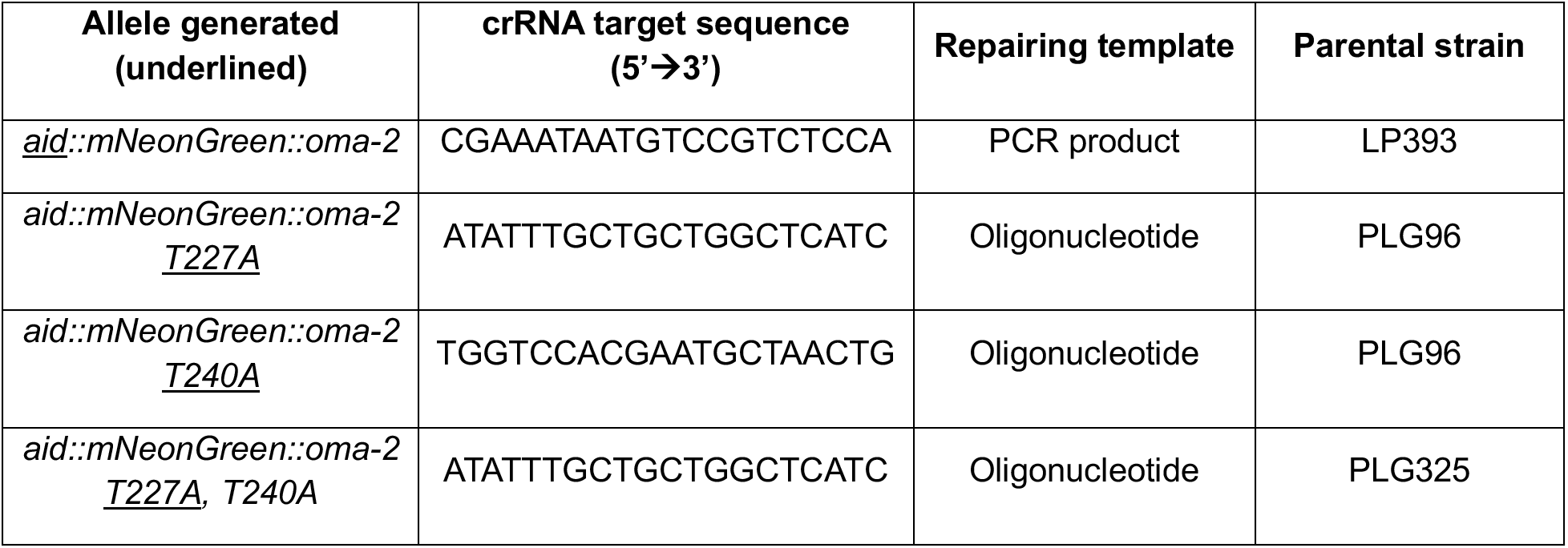
List of CRISPR gRNAs used in this study.

**Table S3:**
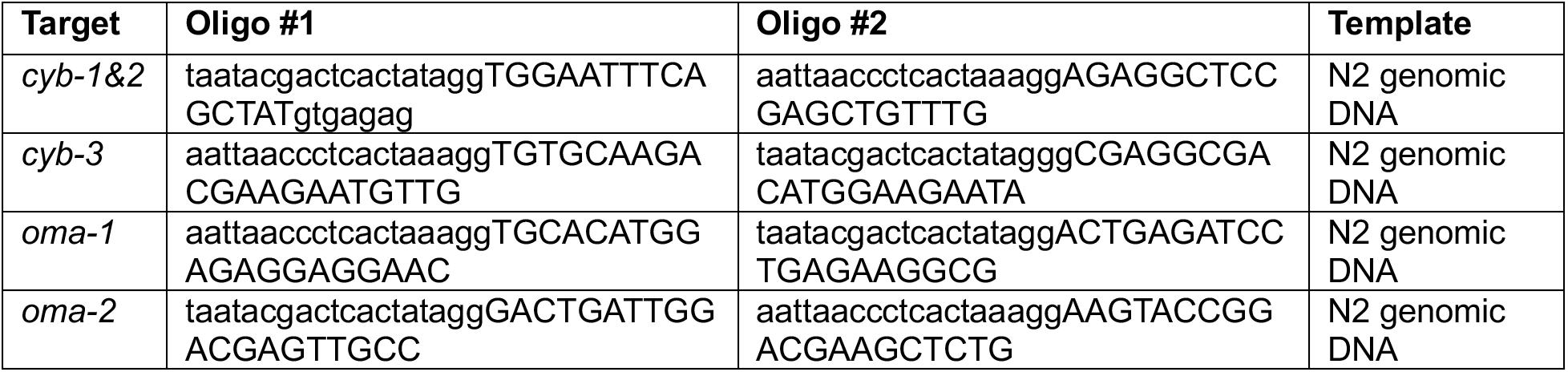
List of oligonucleotides used for dsRNA generation.

**Table S4:** Plasmids used in this study.

| Plasmid | Description |
| --- | --- |
| pLA012 | CDK-1::3xFlag for human cell expression |
| pLA013 | 5xMyc::CYB-3 WT for human cell expression |
| pLA016 | 5xMyc::CYB-3 HP<sup>mut</sup> for human cell expression |
| pLA017 | 5xMyc::CYB-3 PBP<sup>mut</sup> for human cell expression |
| pLA223 | pGEXLB/GST::OMA-2<sup>208-393</sup> for bacteria expression |
| pLA224 | pGEXLB/GST::OMA-2<sup>208-393</sup> T227A for bacteria expression |
| pLA225 | pGEXLB/GST::OMA-2<sup>208-393</sup> T240A for bacteria expression |
| pLA226 | pGEXLB/GST::OMA-2<sup>208-393</sup> T227A, T240A, S291A, T314A, T327A, S335A for bacteria expression |
| pLA227 | CDK-1(D146N)::3xFlag for human cell expression |

## SUPPLEMENTARY MOVIES

**Movie S1.** Time lapse sequences of embryos expressing mCherry-tagged Histone H2b and either OMA-1::GFP or mNeonGreen::OMA-2. Time stamp indicates time elapsed from pronuclear meeting (PNM) in seconds. 5 x 5 µm z-stacks were acquired every 30 seconds and maximum intensity projections generated for each time frame. Playback rate is 10 frames per second, or 300X relative to real time.

**Movie S2.** Time lapse sequences of embryos expressing mCherry-tagged Histone H2b and mNeonGreen::OMA-2; as well as the indicated *cyb-3* transgenes (WT, HP^mut^ or PBP^mut^). Embryos were also depleted of endogenous *cyb-3* through RNAi. Time stamp indicates time elapsed from pronuclear meeting (PNM) in seconds. 5 x 5 µm z-stacks were acquired every 20 seconds and maximum intensity projections generated for each time frame. Playback rate is 10 frames per second, or 200X relative to real time.

